# Building a syntrophic *Pseudomonas putida* consortium with reciprocal substrate processing of lignocellulosic disaccharides

**DOI:** 10.1101/2024.11.19.624300

**Authors:** Barbora Burýšková, Jesús Miró-Bueno, Barbora Popelářová, Barbora Gavendová, Ángel Goñi-Moreno, Pavel Dvořák

## Abstract

Synthetic microbial consortia can leverage their expanded enzymatic reach to tackle biotechnological challenges too complex for single strains, such as lignocellulose valorisation. The benefit of metabolic cooperation comes with a catch – installing stable interactions between consortium members. We constructed a syntrophic consortium of *Pseudomonas putida* strains for lignocellulosic disaccharide processing. Two strains were engineered to hydrolyse and metabolise lignocellulosic sugars: one grows on xylose and hydrolyses cellobiose to produce glucose, while the other grows on glucose and cleaves xylobiose to produce xylose. This specialisation allows each strain to provide essential growth substrate to its partner, establishing a stable mutualistic interaction, which we term reciprocal substrate processing. Key enzymes from *Escherichia coli* (xylose isomerase pathway) and *Thermobifida fusca* (glycoside hydrolases) were introduced into *P. putida* to broaden its carbohydrate utilisation capabilities and arranged in a way to install the strain cross-dependency. A mathematical model of the consortium assisted in predicting the effects of substrate composition, strain ratios, and protein expression levels on population dynamics. Our results demonstrated that modulating extrinsic factors such as substrate concentration can optimise growth and balance fitness disparities between the strains, but achieving this by altering intrinsic factors such as glycoside hydrolase expression levels is much more challenging. This study underscores the potential of synthetic microbial consortia to facilitate the bioconversion of lignocellulosic sugars and offers insights into overcoming the challenges of establishing synthetic microbial cooperation.

## Introduction

In nature, mostly all microorganisms interact on a metabolic basis. This expands the enzymatic toolbox available for transforming their environment. The reach and abundance of microbes make them the geoengineers of new habitats, whose formation drives greater diversity, expanding the chemical conversion toolbox even further. Microorganisms are responsible for running whole biogeochemical cycles on our planet, and they achieve this through diversity and cooperation (Falkowski et al., 2008; Morris et al., 2013).

As synthetic biologists aim to tackle increasingly complex biotechnological tasks, they keep running into the wall of limitations set by tinkering with single-organism monocultures. In the past years, one of the approaches to solving the issue of monoculture limits became the engineering of whole microbial consortia (Brenner et al., 2008; McCarty & Ledesma-Amaro, 2019; Roell et al., 2019; Rapp et al., 2020; Ibrahim et al., 2021; Snoeck et al., 2024). These consist of multiple cooperating organisms, each bearing a part of an envisioned bioprocess to a) complement each other and share the necessary burden (Bokinsky et al., 2011; Minty et al., 2013; Zhou et al., 2015), b) separate incompatible chemical reactions (Shabab et al., 2020), c) support each other by removing waste (Cha et al., 2021), intermediates (Bizukojc et al., 2010), or supplying coenzymes (Cooper et al., 2019). While the engineering limits are theoretically boundless due to the modularity of microbial consortia, establishing stable relationships between the individual consortium members is a recurring problem (Wondraczek et al., 2019; Duncker et al., 2021; Dinh et al.,2020). *De novo* interactions can be installed by quorum-sensing circuits that usually induce negative feedback loops in the consortium strains to maintain community balance (Balagadde et al., 2008; Scott et al., 2017). These, however, cause a high selective force for escapees that mutate or otherwise eliminate the threatening genetic components (Balagadde et al., 2005). Another option is to install mutualistic relationships such as syntrophy or cross-feeding that rely on the consortium members benefiting from the presence of their consortial partners (Rodriguez Amor & Dal Bello, 2019). This approach is more in line with the inherent tendency of organisms to evolve (Castle et al., 2021) and could lead to better long-term stability of the system (D’Souza et al., 2018).

One of the complex tasks tackled by synthetic biology is the valorisation of lignocellulosic residues. The carbon and energy stored in plant waste can be leveraged and upcycled into commodity and specialty chemicals, circularising parts of our economy (Taha et al., 2016), yet, processing this sturdy material is energy-demanding and different approaches to lignocellulose depolymerisation are studied to find an economically feasible option (Kumar et al., 2008; Yamada et al., 2013; Parisutham et al., 2017; Kim et al., 2019). In nature, white- and brown-rot fungi, actinomycetes, and other saprophytic bacteria produce a complex enzymatic cocktail that degrades all three constituents of plant biomass - cellulose, hemicellulose, and lignin (Cragg et al., 2015; Ventorino et al., 2015). This makes lignocellulose valorisation a perfect case for the experimental employment of (synthetic) microbial consortia and the study of syntrophy for the establishment of mutualistic relationships between consortial strains (Das et al., 2023; Minty et al., 2013; Li et al., 2024; Vu et al., 2023).

In this work, we equipped the resilient saprophytic soil bacterium and robust biotechnological chassis *Pseudomonas putida* EM42 (Bugg et al., 2021; Kohlstedt et al., 2022, Martinez-García et al., 2014a,b) with enzymes from *Escherichia coli* (XylA, XylB, XylE) and from the cellulolytic actinomycete *Thermobifida fusca* (BglC, Xyl43A) to broaden its carbohydrate utilisation capacity. We took advantage of a mixed substrate of two lignocellulosic disaccharides (Liu et al., 2020), cellobiose and xylobiose, to divide the heterologous genes between two *P. putida* strains. These were specialised to grow on either xylose (strain CP-X) or glucose (strain CP-G) monomers and cleave xylobiose (strain CP-G) or cellobiose (strain CP-X) disaccharides, respectively, to produce palatable substrate for their partner (**Fig. 1**). This reciprocal substrate processing established a stable mutualistic relationship. Accompanied by a mathematical model describing the consortium behaviour, we further explored how modulating substrate composition, strain ratios, and protein expression can affect the population dynamics of our system. Our syntrophic consortium of two specialised strains divides the carried burden (Snoeck et al., 2024) and circumvents the potential issue of carbon catabolite repression (CCR; Görke & Stülke, 2008; Rojo, 2010). On the other hand, differences in strain fitness lead to unequal growth rates. This demonstrates that each synthetic consortium approach has its challenges and the substrate, the strain build, and the cultivation conditions all play a role in the optimisation of the process, making consortia a great opportunity for Lego-like combinatorics, application of new molecular tools, and encountering (and perhaps answering) many basic research questions along the way.

**Figure 1.**
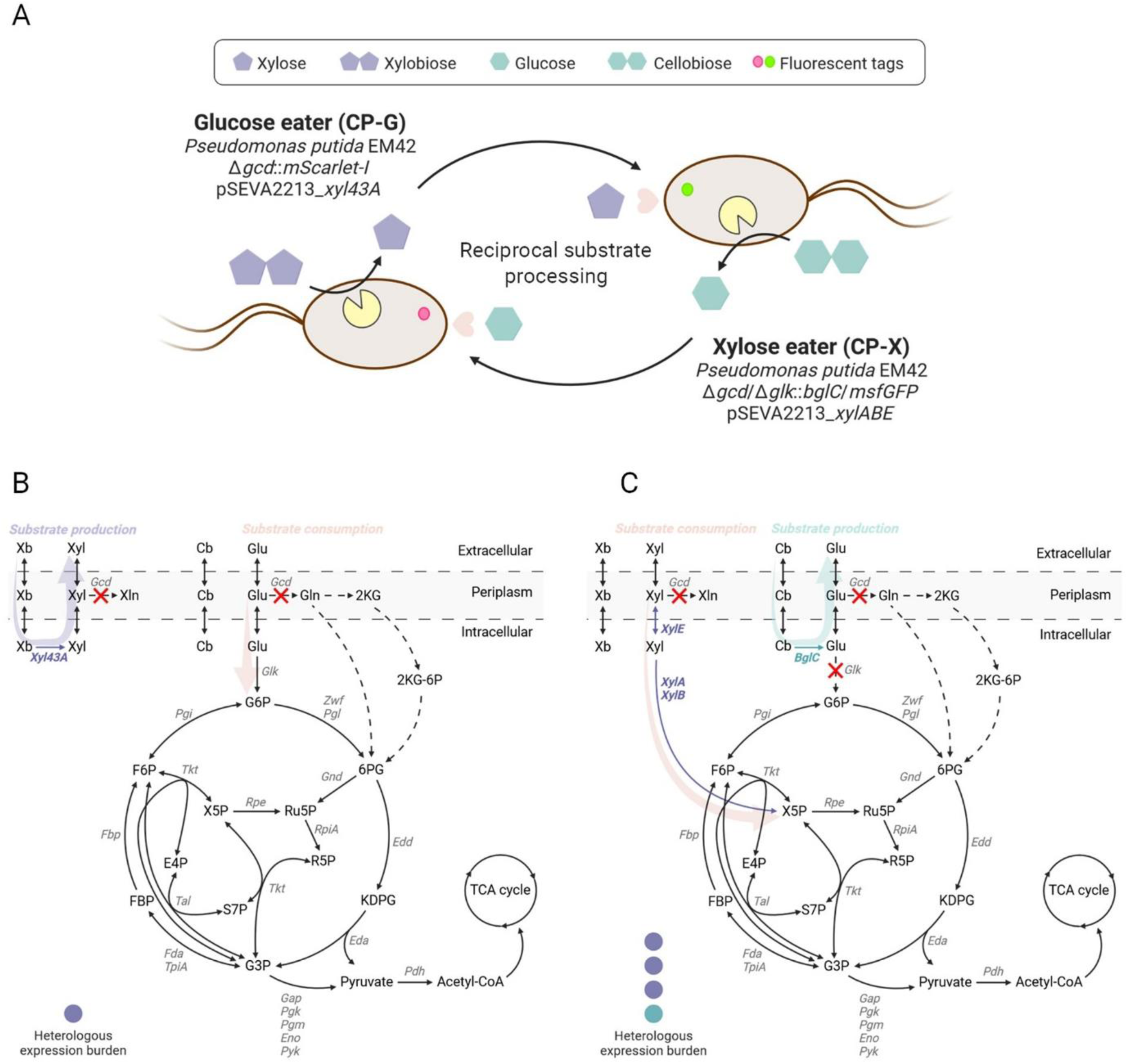
Consortium blueprint (**A**) and schematic illustration of sugar metabolism in the constructed *P. putida* strains CP-G (**B**) and CP-X (**C**). Heterologous enzymes and pathways are depicted in purple and teal. Deleted reactions are marked by a red cross. Native but vacant pathways are depicted by dotted arrows. The expected level of metabolic burden placed on the strains by heterologous expression is shown. Enzyme abbreviations: BglC β-glucosidase, Eda 2-keto-3-deoxy-6-phosphogluconate aldolase, Edd 6-phosphogluconate dehydratase, Eno phosphopyruvate hydratase, Fbp fructose-1,6-bisphosphatase, Fda fructose-1,6-bisphosphate aldolase, Gap glyceraldehyde-3-phosphate dehydrogenase, Gcd glucose dehydrogenase, Glk glucokinase, Gnd 6-phosphogluconate dehydrogenase, Pdh pyruvate dehydrogenase, Pgk phosphoglycerate kinase, Pgm phosphoglycerate mutase, Pyk pyruvate kinase, Rpe ribulose-5-phosphate 3-epimerase, RpiA ribose-5-phosphate isomerase, Tal transaldolase, Tkt transketolase, TpiA triosephosphate isomerase, XylA xylose isomerase, XylB xylulokinase, XylE xylose symporter, Xyl43A β-xylosidase, Zwf glucose-6-phosphate dehydrogenase. Metabolite abbreviations: Cb cellobiose, E4P erythrose 4-phosphate, FBP fructose 1,6-bisphosphate, F6P fructose 6-phosphate, Gln gluconate, Glu glucose, G3P glyceraldehyde 3-phosphate, G6P glucose 6-phosphate, KDPG 2-keto-3-deoxy-6-phosphogluconate, 2KG 2-ketogluconate, 2KG-6P 2-ketogluconate 6-phosphate, 6PG 6-phosphogluconate, R5P ribose 5-phosphate, Ru5P ribulose 5-phosphate, S7P sedoheptulose 7-phosphate, Xb xylobiose, Xln xylonate, X5P xylulose 5-phosphate.

## Results and discussion

### Design of the consortium

The design of our consortium was guided by a few key principles. We wanted each strain to specialize in utilizing one type of sugar substrate - glucose for strain CP-G and the *P. putida* non-native substrate xylose for strain CP-X. Such an approach prevents potential sequential substrate uptake or crosstalk caused by CCR (Rojo, 2010; Flores et al., 2019), and it allowed us to build two independent populations as the basis for our consortium. We chose the disaccharide versions of each sugar - cellobiose and xylobiose - as the primary substrates, since oligomer degradation represents a step towards consolidated bioprocessing of lignocellulose, allowing for cheaper pretreatment of plant waste (Chen, 2015; Yamada et al., 2017). The other reason was that heterologous expression of the glycoside hydrolases needed for cleaving the two disaccharides presented us with a unique opportunity to set up a mutual relationship between the two populations. Each strain would express the glycoside hydrolase cleaving the substrate to be assimilated by the second strain. This reciprocal substrate processing made both strains dependent on one another in a form of syntrophy (**Fig. 1**; Morris et al., 2013; D’Souza et al., 2018).

Substrate specificity was ensured by the lack of a xylose utilisation pathway in wild-type *P. putida* which became the Glucose eater (CP-G). Heterologous plasmid-based expression of *xylABE* genes from *E. coli* encoding the xylose isomerase pathway and xylose transporter (pSEVA2213_*xylABE*; Dvořák et al., 2018) was the basis of the Xylose eater (CP-X) together with a deletion of the glucokinase gene *glk* (PP_1011) disabling the growth on glucose. Both strains harboured a deletion of the periplasmic glucose dehydrogenase gene *gcd* (PP_1444) to block the conversion of xylose to the dead-end product xylonate (Dvořák et.al, 2018; **Fig. 1**). To distinguish the strains, fluorescent tags preceded by a translational coupler were introduced into the chromosome via the mini-Tn7 transposon system from Zobel & Benedetti, 2015, targeting the genes into the *att*Tn7 site for stable basal expression. The Xylose eater was tagged with msfGFP while the Glucose eater was tagged with mScarlet-I (Schlechter et al., 2015). The reciprocal substrate processing was established by the heterologous expression of glycoside hydrolases from the cellulolytic actinomycete *Thermobifida fusca*. The Glucose eater produced Xyl43A β-xylosidase (Tfu_1616) from pSEVA2213_*xyl43A*, whereas the Xylose eater produced BglC β-glucosidase (Tfu_0937), whose gene was genome-integrated via the mini-Tn5 transposon system (Martínez-García & de Lorenzo, 2012). Both enzymes were expressed with an N’-terminal His tag and should be located intracellularly (Moraïs et al., 2012; Spiridonov & Wilson, 2001). The setup of the strains was expected to place a higher metabolic burden on the Xylose eater due to the load of heterologous enzymes expressed in addition to metabolising a non-native substrate. Nevertheless, the reciprocal substrate processing was expected to establish cooperative dependency.

### Components testing and initial consortium characterisation

The consortium members held their respective substrate specificities. When first constructed, the Xylose eater showed limited growth in liquid medium with its respective substrate xylose (**Supplementary Fig. 1**). Since the phenotype was confirmed on solid medium (not shown), we decided to culture the strain in liquid medium with xylose until growth commenced. We then inoculated fresh xylose medium twice and the time needed for the culture to reach visible growth plummeted from six to three to two days. The initial growth stagnation was initially thought to be caused by the *glk* deletion affecting the regulation of neighbouring genes encoding enzymes of the EDEMP cycle (Daddaoua et al., 2009). But after the template (CP-Xpre) and improved strain (CP-X) were sequenced, a large genome multiplication (275 kb) was found in CP-X encompassing 236 genes in two or three copies framed by IS4 family transposase ISPpu8 (**Supplementary Table 4**). This multiplication included transaldolase *tal* [PP_2168] involved in the pentose phosphate pathway (PPP) which is strongly engaged in xylose utilisation, and glyceraldehyde-3-phosphate dehydrogenase *gapB* [PP_2149] funnelling carbon out of the EDEMP cycle towards the TCA cycle. Multiplication of the same genetic region occurred in the adaptive laboratory evolution of *P. putida* EM42 on xylose in the study by Dvořák et al., 2024, where the effect of *tal* overexpression on xylose utilisation was confirmed by reverse engineering. The reoccurrence of this adaptive genome rearrangement shows how quickly *P. putida* can adjust its metabolic capacity to different directions of carbon influx.

In both consortium strains the glycoside hydrolases - β-glucosidase BglC and β-xylosidase Xyl43A - were well expressed. BglC made up to 12,3 % of the total protein in CP-X and CP-G contained 26 % of Xyl43A as visualised by SDS-PAGE (**Supplementary Fig. 2**). We then tested their activity by observing disaccharide cleavage in resting cells using HPLC analysis (**Fig. 2**). We confirmed that the intracellular hydrolytic activity of both enzymes, together with minor background activity of molecules that leached out of the cells, efficiently produces monosaccharides. Both strains are thus capable of supplying substrate for their consortial counterpart. Since the majority of enzymatic activity is intracellular, this suggests that a transport mechanism for cellobiose and xylobiose is present in *P. putida* EM42 (**Fig. 2**). The transport of cellobiose into the cytoplasm of *P. putida* has already been reported, unlike the transport of the non-native disaccharide xylobiose (Dvořák & De Lorenzo, 2018). The effect of transport mechanisms on microbial metabolism and ensuing bioengineering efforts are an understudied and thus an interesting topic for further exploration (Van der Hoek & Borodina, 2020).

**Figure 2.**
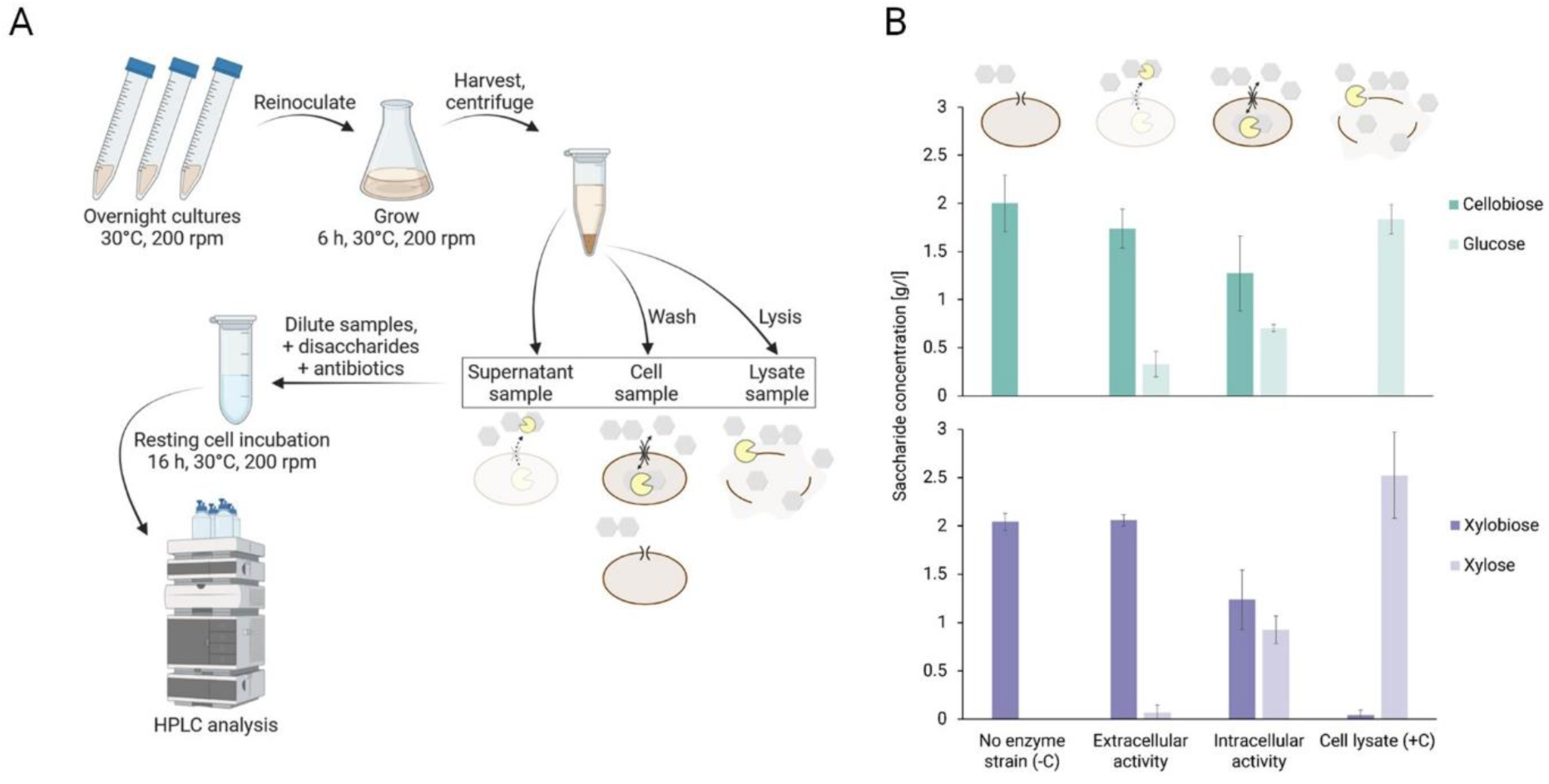
Resting cell experiment verifying disaccharide cleavage by heterologous glycoside hydrolases and transport through the cellular envelope. **A**) Workflow of the experiment. Details can be found in the Methods section. **B**) Conversion of disaccharides to monosaccharides by consortium strain cells (Intracellular activity), their culture supernatants (Extracellular activity), or lysates (Cell lysate +C), and negative controls (No enzyme strain -C, ID18 and ID24) after 16 hours of incubation with 2 g/l of disaccharides. BglC activity is depicted in the upper graph, Xyl43A activity is depicted in the lower graph. Columns represent means ± standard deviations calculated from four (n = 4) biological replicates from two independent experiments.

Basal fluorescent protein expression enabled easy strain distinction under the microscope as well as during growth experiments in plate reader format (**Fig. 3**). We compared continual calibration during growth on monosaccharides with calibration using LB-grown overnight cultures and decided to use continual calibration on monosaccharides throughout the project since it better reflected the OD (**Supplementary Fig. 3**). The slower maturation of mScarlet expressed by CP-G led to calibration bias and while it served well in reporting the general state of strain ratios, we decided to use the measured OD of all cells subtracted by the reliable GFP signal of CP-X cells for graphical depiction of CP-G cells in cocultivations.

**Figure 3.**
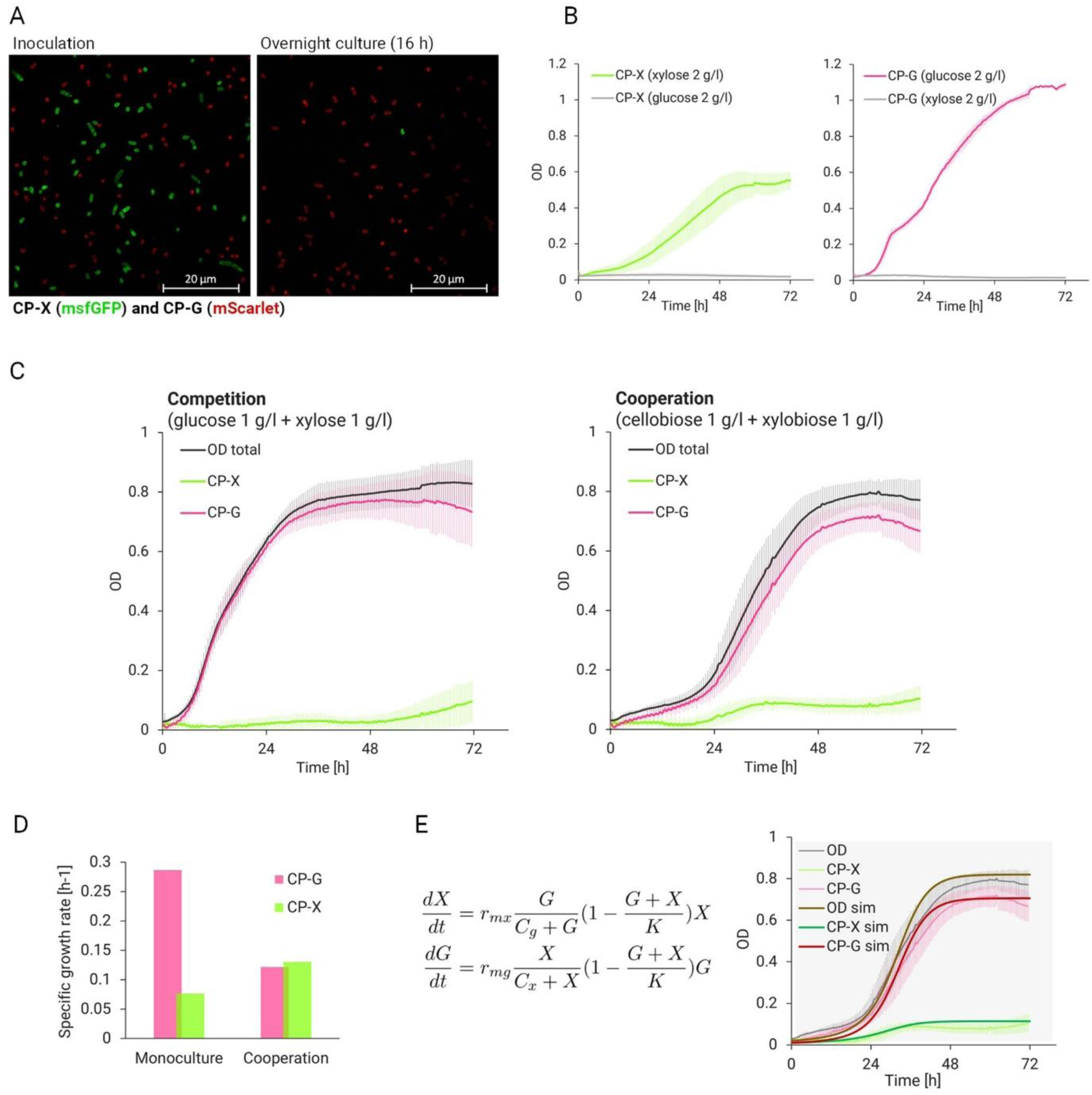
Characterisation of consortium strains. (**A**) Micrographs of basal fluorescence recorded with ZEISS Elyra 7. CP-G tagged by mScarlet and CP-X tagged by msfGFP in a 1:1 strain ratio at the beginning (left) and after overnight cultivation (16 h, right) of the consortium in complex LB medium. (**B**) Monoculture growth in 96-well plate format on respective monosaccharides. Data are shown as means ± standard deviations from several independent experiments and multiple biological replicates (CP-X on xylose n = 13, CP-X on glucose n = 9; CP-G on glucose n = 7, CP-G on xylose n = 8). (**C**) Cocultivation in 96-well plate format on monosaccharides (competition; left) and on disaccharides (cooperation; right). Data are shown as means ± standard deviations from nine independent experiments and 18 (n = 18) biological replicates. In (**D**) the changes in specific growth rate between monocultures (from B) and cooperation (C, right) are shown. (**E**) Mathematical model of the consortium. The behaviour of the consortium is described by two differential equations (left). The variables *X* and *G* represent the optical density (OD) of the strains CP-X and CP-G, respectively. The parameters *r_mx_* and *r_mg_* are the *maximum per capita growth rates* for *X* and *G*. The factor *G/(C_g_+G)* represents the transformation of xylobiose into xylose by strain CP-G, while the factor *X/(C_x_+X)* represents the transformation of cellobiose into glucose by strain CP-X. The parameters *C_x_* and *C_g_* account for the amount of glycoside hydrolases. Specifically, *C_x_* and *C_g_* are inversely proportional to the BglC and Xyl43A enzymes, respectively. The factor *(1 - (G + X) / K)* indicates the fraction of the free carrying capacity of the medium, where parameter *K*, known as the *carrying capacity*, depends on available resources. Data gathered from cooperation on disaccharide experiments was used to build a mathematical model describing the behaviour of the cooperating consortium (right). The initial conditions are *X_0_*=0.01 and *G_0_*=0.01.

Both members of the consortium were then cocultivated on monosaccharides (1 g/l xylose + 1 g/l glucose) and disaccharides (1 g/l xylobiose + 1 g/l cellobiose). The different cultivation conditions enabled the observation of competitive behaviour on monosaccharides and cooperative dependency on disaccharides (**Fig. 3C**). On monosaccharides, no carbon cross-feeding was needed and thus the growth rate differences dictated the outcome where strain CP-G overgrew strain CP-X. On the disaccharides, the dependency of the two strains was indicated by the long lag phase of the whole consortium (∼20h) corresponding to the lag phase of CP-X alone, and by the simultaneous transition of both strains into the exponential phase. Since the dependence is enabled by the activity of the heterologously expressed glycoside hydrolases the onset of the delayed exponential phase of CP-G was thus thought to mark the accumulation of enzyme and resulting monosaccharide in the culture. The established relationship also altered the maximal growth rate of the individual strains when grown in a cooperating consortium compared to monocultures grown on monosaccharides. Whereas the growth rate of CP-X benefited from the cooperation, the growth rate of CP-G fell by over 60% (**Fig. 3D**). This shows that although the growth of the monocultures is not comparable, the engineered relationship establishes dependence of both strains in reciprocal substrate supply which affects the growth pattern of the consortium members.

### Mathematical modelling of consortium behaviour

The gathered cocultivation data was used to build a mathematical model describing the behaviour of the consortium when grown on disaccharides. The model consists of two differential equations that describe the logistic growth of each strain (**Fig. 3E**). This modelling approach allows us to represent the cross-feeding interactions between the two bacterial strains. The model builds on established principles in microbial dynamics which explore logistic growth models for continuous population dynamics (Murray, 2002; Balagaddé et al., 2008; Freilich et al., 2011; Tsoi et al., 2018).

The accuracy of our model was validated by predicting the growth of the consortium strains cocultivated on disaccharides and comparing it to the measured data (**Fig. 3E**). We manually fitted the parameters *r_mx_*, *r_mg_*, *C_x_*, *C_g_*, and *K* of the model using the total OD data from the experiment and the CP-X OD data from the calibration. Then, we plotted the OD of the strain CP-G and checked that the predicted OD of CP-G was similar to the one obtained using the equation OD_total_ = OD_x_ + OD_g_. The model was then employed to search for cultivation variables (substrate ratios, inoculation ratios), that, when adjusted, would balance the growth of the two consortial strains. To make predictions for cultivation variables, we used scaling factors represented by the symbol *α*. Specifically, *G/(C_g_+G)* is scaled based on the initial amount of xylobiose (*α_Dg_*), and *X/(C_x_+X)* is scaled based on the initial amount of cellobiose (*α_Dx_*). The parameter *r_mx_* is scaled by the burden due to BglC expression and degradation (*α_Bx_*). The parameter *C_x_* is scaled according to the amount of BglC (*α_Ex_*). Additionally, the value of *K* is scaled by both the initial amount of disaccharides and the amount of the BglC enzyme, represented by *α_K_*, where *α_K_ = min{α_Ex_, α_Dx_, α_Dg_}*. With these adjustments, the final form of the two differential equations shown in **Fig. 3E** is:

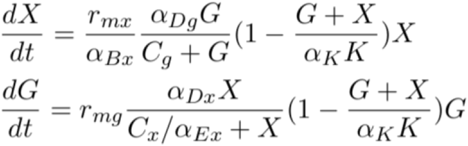

For the consortium experiment in **Fig. 3E** the scaling factors *α_Dx_*, *α_Dg_*, *α_Bx_*, and *α_Ex_* were all set to 1 and the parameter values were *r_mx_*=0.12 1/h, *r_mg_*=0.34 1/h, *C_x_*=0.04, *C_y_*=0.02 and *K*=0.82. The benefit of the mathematical model developed in this study is its ability to explore the system’s potential beyond what is immediately observable in the lab. The process began with experimental work, followed by calibrating the model based on these experiments. This calibrated model then allowed us to conduct *in silico* studies to explore the various possibilities within the system, such as optimizing cultivation conditions and predicting system behaviour under different scenarios.

### Mathematical model enables searching for optimal cultivation conditions

A possible means of balancing the consortium when strain growth rates are unequal is choosing inoculation ratios that favour the slower growing strain (Dinh et al., 2020). We thus repeated the cocultivation with inoculation ratios 2:1, 4:1, and 10:1 in favour of CP-X. None of the cultivations favouring CP-X balanced the strain growth curves (**Fig. 4A**). Additionally, a higher inoculation ratio in favour of CP-X only prolonged the lag phase of both strains. The CP-G overgrowth trend stayed and shows that the relationship is stable, as observed for other obligate cross-feeding consortia (Losoi et al., 2019). Since an advantage in numbers did not help CP-X to achieve better growth, as predicted by the model (**Fig. 4C**), we tried to cultivate the consortium on different ratios of disaccharides which could give us more insight into the consortium dynamics and another opportunity to challenge the predictions of our mathematical model.

**Figure 4.**
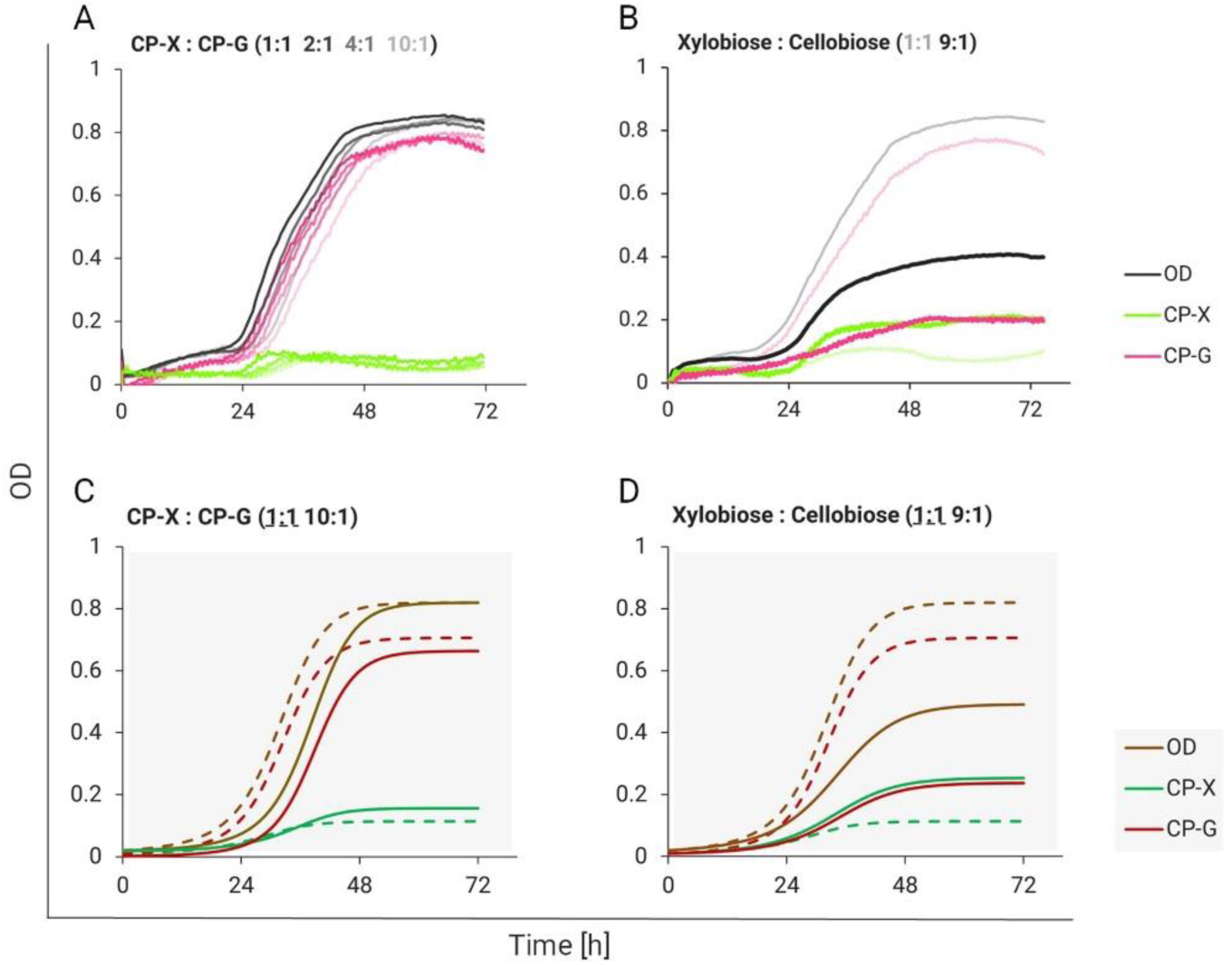
Consortium cocultivation on disaccharides (1+1 g/l) with different strain inoculation ratios (**A**) and substrate ratios (**B**) in 96-well plate format. Data are shown as means from at two (n = 2) biological replicates. Error bars are omitted for clarity. The lower two graphs show respective model-predicted outcomes. (**C**) Simulation of the consortium with initial conditions *X_0_*=0.0182 and *G_0_*=0.0018. (**D**) Simulation of the consortium with high xylobiose (*α_Dg_*=1.4) and low cellobiose (*α_Dx_*=0.6).

Altering the substrate ratio balanced the growth of our consortium strains. When limiting the Glucose eater with substrate (9 parts xylobiose : 1 part cellobiose), the resulting ratio of CP-X and CP-G in the consortium was 1:1 (**Fig. 4B**). However, the final OD of the consortium was two times lower compared to the consortium grown on equal amounts of both disaccharides. Taken together, our data show in line with the mathematical prediction (**Fig. 4D**) that while inoculation ratios do not balance the growth of the cocultivated strains, the substrate ratios do, which could be leveraged in a continuous cultivation system (Govindaswamy & Vane, 2010). On the other hand, asymmetric growth can be leveraged by adding an anabolic pathway to the faster growing strain (CP-G). This would introduce a production module to the system while balancing the burden placed on both consortium members - turn your challenges into opportunities.

### Searching for bottlenecks

To see how fast the saccharide production and consumption are in real-time, we repeated the cultivation of the consortium and individual consortium strains on monosaccharides and disaccharides and included regular sampling for HPLC analysis (**Fig. 5**).

**Figure 5.**
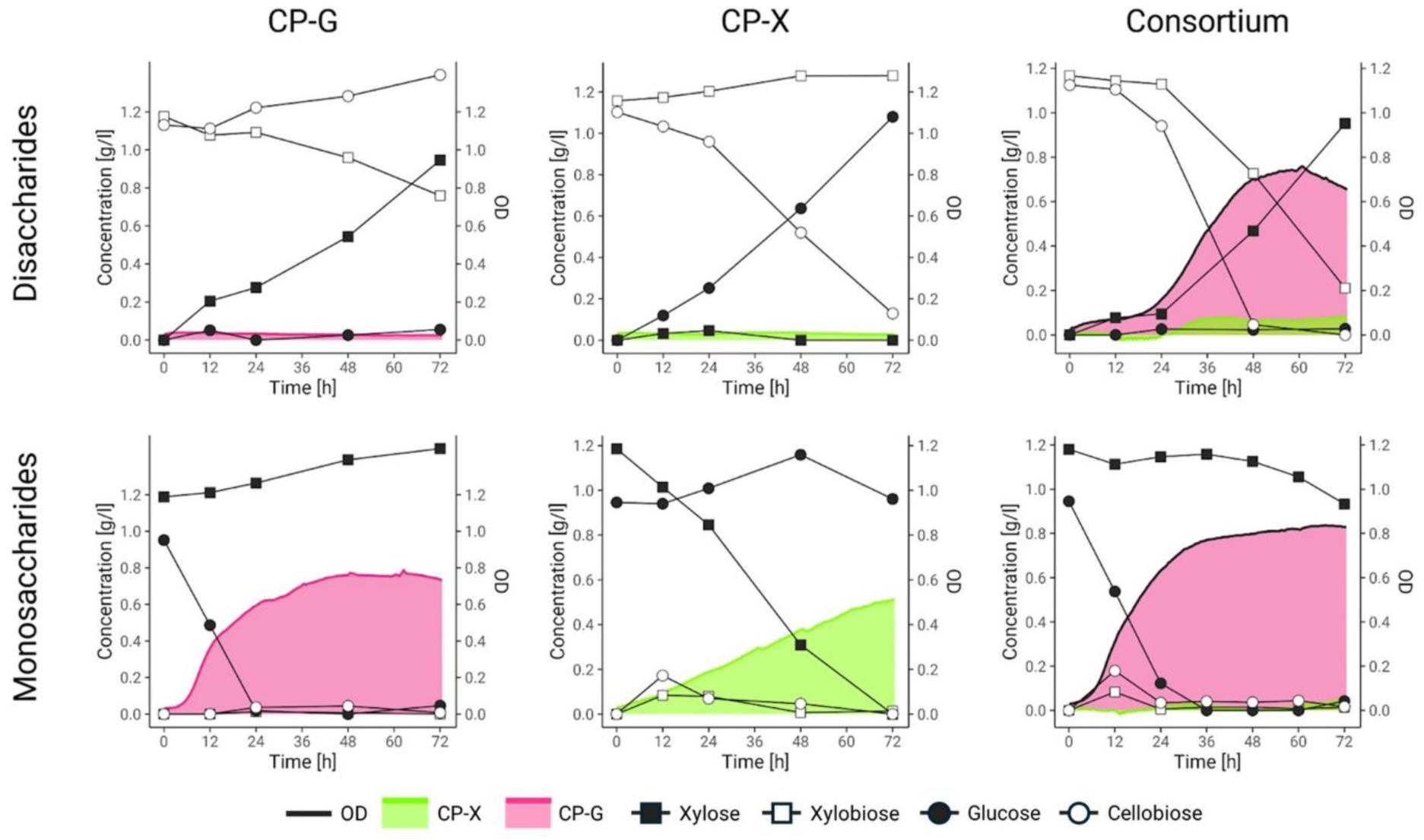
Consortium and monoculture cultivations on disaccharides and monosaccharides in 96-well plate format with saccharide concentration monitoring by HPLC. OD is depicted as an area mapped on the right y-axis, whereas sugar concentrations are depicted with line graphs mapped on the left y-axis. Data are shown as means from at least three (n ≥ 3) biological replicates from two separate experiments. Note that slight evaporation of culture volume towards the end of the experiment causes sample concentration, which makes the amount of abundant sugars seem to rise.

The experiment was carried out in a 96-well plate format for direct comparison with earlier cultivations. Multiple technical replicates were used to ensure sufficient sampling volumes. As expected, CP-X and CP-G individually only consumed their respective monosaccharide substrates. Interestingly, the growth patterns slightly differed from ones obtained on single monosaccharides (**Fig. 3B**), pointing towards altered sugar assimilation regulation when two types of sugar substrates are present. When grown individually on disaccharides (**Fig. 5**), no growth occurred, yet, the inoculation biomass itself (250 µl of OD 0,05) contained enough glycoside hydrolases to convert the majority of the respective disaccharide to monosaccharide. This confirmed that the enzymes are well expressed (**Supplementary Fig. 2**) and active, and should not pose a bottleneck in strain cross-feeding. Cocultivation of the consortium strains on monosaccharides showed the rapid consumption of glucose by the CP-G strain and a minimal decrease in xylose consumed by the negligible growth of the CP-X strain (**Fig. 5**). A temporary rise of disaccharides (around 12 h) is possibly caused by the reverse activity of the glycoside hydrolase enzymes in the presence of a high product-to-substrate ratio (Bhatia et al., 2002). Cocultivation on disaccharides shows that even though both glycoside hydrolases are active, conversion of cellobiose to glucose is faster when both strains enter exponential phase (Consortium -Disaccharides; see incline in cellobiose and xylobiose concentration between 24 h and 48 h). At the end of the experiment, the majority of xylose is still available in the growth medium. The fitter CP-G strain probably depletes other micronutrients or oxygen in the medium, disabling further growth of CP-X.

### Tightening the mutualistic dependence with degradation tags

After gathering the information from the first Design-Build-Test-Learn cycle, for the second round we decided to tighten the consortium members’ relationship. The major molecular parts that enable the mutualistic relationship are the glycoside hydrolases, and from the HPLC analysis of the cocultivation medium we knew, that β-glucosidase produced by CP-X hydrolysed disaccharides faster than β-xylosidase from CP-G (**Fig. 5**, Consortium, Disaccharides). We thus decided to link β-glucosidase (BglC) activity more tightly to the growth of CP-X by increasing its degradation rate to prevent its accumulation. This would limit the rate at which glucose substrate for the CP-G strain would be produced. A simulation by the mathematical model supported our idea, that decreasing the amount of BglC could help establish a more balanced growth of the consortial strains (**Fig. 6A**). For this, we introduced sequences to the 3′-end of the *bglC* gene that would translate into C-terminal degrons and mimic natural proteolytic signalling recognized by the native ClpXP protease complex (Izert et al., 2021). We chose a *Pseudomonas*-specific SsrA degradation tag (AANDENYALAA) together with its two-amino acid substitution variant used as a control in Keiler et al., 1996 (AANDENYALDD; Keiler et al., 1996; Karzai et al., 2000; Durante-Rodríguez et al., 2020) and a poly-A degradation tag (AAAAAA), representing a conserved ribosome-associated quality control mechanism mediated by RcqH (Lytvynenko et al., 2019; **Fig. 6B**). The tags were introduced in front of the STOP codon of the *bglC* gene in the chromosome of strain CP-X using pSNW2-based integration vectors constructed by *in-vivo* cloning (Wirth et al., 2020; **Fig. 6C**).

**Figure 6.**
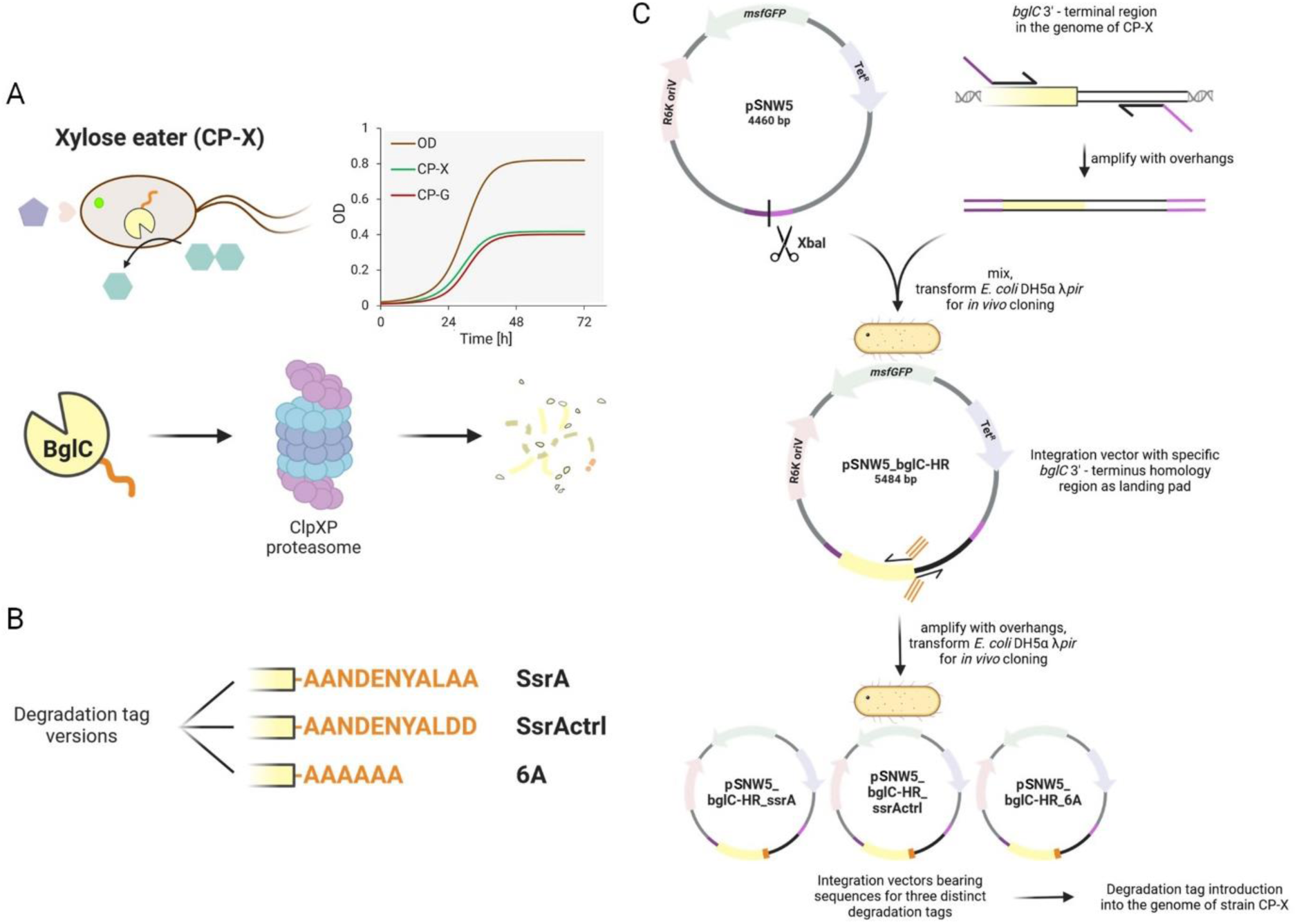
Reduction of BglC excess in CP-X by degradation tags. **A**) Schematic illustration of the degradation-tagged BglC fate with a simulation of the expected effect on the consortium dynamics (for the simulation, *C_x_* was set to 0.08 and *r_mx_* to 0.22 h^-1^). **B**) Amino acid sequences of the three degradation tags. **C**) Cloning workflow for targeted addition of the degradation tags into the CP-X genome. Details can be found in the Methods section.

The CP-X strain versions with degradation-tagged BglC were first grown individually to test their growth (**Fig. 7**). All degradation-tagged BglC strains grew worse than the template strain. We argue that this was caused by the highly efficient targeting of their β-glucosidase (12,3 % of cellular protein; **Supplementary Fig. 2**) to ATP-dependent proteasomes. We confirmed this by SDS-PAGE analysis, which showed that in the CP-X_ssrA strain all BglC was degraded (0 % of cellular protein; **Supplementary Fig. 2**) and in CP-X_6A 75 % of it was degraded (3,1 % of cellular protein; **Supplementary Fig. 2**). Yet, why the growth of CP-X_ssrActrl was impaired is not clear (11,5 % of cellular protein; **Supplementary Fig. 2**). According to Fritze et al., 2020, SspB, a mediator protein involved in SsrA-based protein degradation, could still recognise the amino acid sequence of the _ssrActrl control tag. This bond could make SspB unavailable for the native functions of the ribosome rescue pathway and protein recycling and thus slow down cellular growth even without costly BglC degradation.

**Figure 7.**
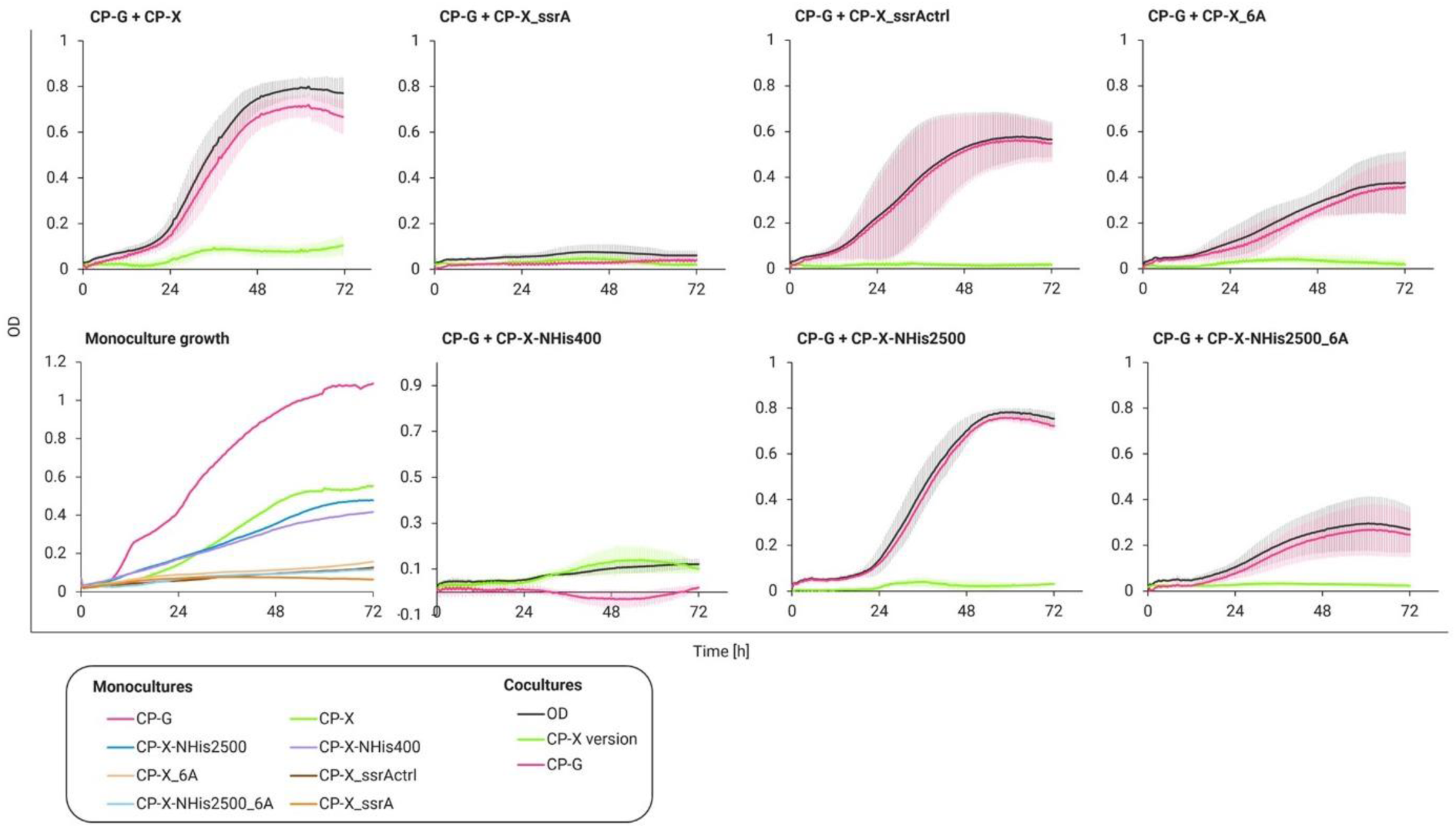
Consortium cooperation in 96-well plate format on disaccharides (1+1 g/l) with CP-X strain variants representing different BglC expression and degradation patterns, leading to modulated relationships of both consortial strains. Monoculture growth on xylose (2 g/l; CP-X) or glucose (2 g/l; CP-G) shows individual strain burden (bottom left) that affects the consortium growth as well. Experimental data are shown as means ± standard deviations from at least three (n ≥ 3) biological replicates from two independent experiments.

Next, the strains were cocultivated as consortia on disaccharides and consortia with BglC degradation were compared to the template consortium (**Fig. 7**). The high BglC degradation activity of the ssrA tag completely disabled the growth of CP-G, and of the whole consortium. Nevertheless, the relationship and thus the fate of CP-G and CP-X strains were tightly linked in this consortium. Microscopic imaging revealed an interesting finding, as this consortium formed clusters of cells after cocultivation (**Supplementary Fig. 4**). To test whether this effect was caused by the strong mutual dependency in which proximity of the cells could be beneficial (Ebrahimi et al., 2019), stress as a variable had to be taken out of the equation since it causes cell clumping as well. In *P. putida* the stress-response sigma factor σRpoS activates the expression of adhesin LapF involved in cell-cell interaction (Martínez-Gil et al., 2010; Lahesaare et al., 2014). The consortium was cultured in pure M9 medium without a carbon source as a stress control. Both the experiment and control sample were cultured in 96-well plate format for 24 hours for microscopy imaging where a 5 x 5 grid of 60 x 60 μm pictures was taken and the cell clusters of this area were counted. The clump ratio of CP-X only : mixed : CP-G only was 1:2:1 in the control sample and 1:3:1 in the experiment sample (**Supplementary Fig. 4**). Although a larger dataset should be examined, this preliminary experiment suggests that the cells are actively attaching to their cognate strain and spatially organising as a response to the engineered relationship (Pignon et al., 2024).

Both the 6A and ssrActrl degradation tags resulted in a lower final OD of the consortium, with most biomass still belonging to CP-G (**Fig. 7**). The lag phase differed in length from the template consortium, perhaps due to stress and cell lysis of CP-X enabling more BglC to escape into the medium and produce glucose faster. Overall, the effect of both the ssrA and 6A degradation tags was confirmed, yet none of the variants led to viable tightening of the strain growth curves. We hypothesised that the high production (RBS strength of 13,018 a.u. leading to 12,3% of total cellular protein in SDS-PAGE **Supplementary Fig. 2**) and subsequent degradation of BglC leads to an unnecessary burden on the Xylose eater and *bglC* expression must be tuned down before the addition of degradation tags for these to have any positive effect.

We thus constructed CP-X variants with weakened expression of *bglC*. Firstly we only removed the N-terminal histidine tag resulting in strain CP-X-NHis2500 with a *bglC* RBS strength of 2,589 a.u. Secondly, we removed the N-terminal histidine tag and additionally switched the original RBS to an *in silico*-designed RBS sequence (Salis Lab RBS calculator - design mode; Salis et al., 2009) for an even lower RBS strength of 417 a.u. resulting in strain CP-X-NHis400. These two CP-X strains with lowered BglC expression resulted in very different consortium outcomes. In the case of the CP-X-NHis2500, the consortium grew almost identically to the template consortium, pointing to a persisting excess of BglC enzyme. On the other hand, strain CP-X-NHis400 did not allow the consortium to grow at all. The RBS strength reduction in strain CP-X-NHis400 was considerable, resulting in no BglC expression (0 % of cellular protein; **Supplementary Fig. 2**), which explains the nonexistent consortium growth. We also constructed one strain in which we combined the two approaches by lowering BglC expression in the CP-X_6A resulting in CP-X-NHis2500_6A. This combination still led to a high burden placed on the xylose eater and the consortium grew to an even lower OD than the CP-X_6A strain consortium.

These outcomes have shown us that finetuning the glycoside hydrolase expression is an easy way to modulate the relationship of our consortium. We did however not find the sweet spot which would allow both strains to grow at a similar rate while reaching a high OD. It is possible that exploring more of the design space (Gilman et al., 2021) would lead to the envisioned outcome. Employing the developed mathematical model, that successfully simulated the consortium behaviour with all CP-X variants (**Supplementary Fig. 5**), to make predictions in the right direction is another possibility. In hindsight, we recognise degradation tags to be a very good tool, but not the best one for a low-burden fix of unbalanced growth rates of consortium strains. Finetuning the relationship bolts expression (glycoside hydrolases in our case) or timing the expression by placing the genes into naturally occurring operons with desired expression characteristics could be an economical way to modulate the population dynamics. Alternatively, an anabolic pathway placed in the fast-growing CP-G strain could have the same effect with added functionality.

## Conclusions

In this study, we engineered two *P. putida* strains for reciprocal substrate processing of two lignocellulosic disaccharides - cellobiose and xylobiose - to explore how little steps towards consolidated bioprocessing can give opportunities for establishing mutualism in synthetic microbial consortia. We divided the catabolic routes for each sugar type - pentose (xylose, metabolised by strain CP-X) and hexose (glucose, metabolised by strain CP-G) - but placed the cognate glycoside hydrolases in the opposite strains to create dependencies that would lead to an obligate mutualistic relationship (Morris et al., 2013; D’Souza et al., 2018). The glycoside hydrolases (BglC and Xyl43A) represented the bolts of this relationship and adjusting their expression or turnover (via degradation tags) was an easy way to modulate the population dynamics of the two consortium strains. Throughout this exploration, we confirmed how much burden is placed on cells through genetic engineering and that for any envisioned bioprocess the “energetic price” of circuits and pathways must be kept to a minimum (Snoeck et al., 2024). Besides modulating the population dynamics of the system intrinsically (through genetic parts), extrinsic factors like substrate ratios or inoculation ratios can also lead to desired changes in the system. The mathematical model, based on differential equations representing the logistic growth of each strain, allowed us to explore the effects of these extrinsic factors. By accurately predicting growth outcomes, the model confirmed that modulating substrate ratios was an effective strategy for balancing the growth of the two strains in coculture. The differences in growth rates could also be leveraged for the addition of new functions to the less-burdened strain. The modular nature of synthetic consortia gives researchers multiple degrees of freedom in designing the catabolic and anabolic pathways, but also the biotechnological process itself (Said & Or, 2017; Said et al., 2020). The advantages of modularity and complexity, however, come with their own set of challenges. In our synthetic consortium, we successfully established a stable mutualistic relationship through reciprocal substrate processing. The population balance was however tilted, as dictated by strain burden. We managed to level this balance by modulating extrinsic factors (substrate concentration) but not through intrinsic factors, where finding a sweet spot in the design space of key genetic components (in this case exogenous glycoside hydrolases) was needed.

## Material and methods

### Cultivation conditions

All strains used and prepared in this study are listed in **Supplementary Table 1**. For plasmid and strain construction, *E. coli* and *P. putida* strains were grown in lysogeny broth (LB) supplemented with appropriate antibiotics at 30 °C and 37 °C, respectively. For growth experiments, night cultures were grown in LB with antibiotics. These were centrifuged (all viable cells were centrifuged at 1,500 g) and resuspended in M9 minimal medium, which consists of M9 salts (7 g/l Na_2_HPO_4_·7H_2_O, 3 g/l KH_2_PO_4_, 0.5 g/l NaCl and 1 g/l NH_4_Cl; pH 7.2), 2 mM MgSO_4_, 100 μM CaCl_2_, 3 ml/l trace element solution (0,3 g/l H_3_BO_3_, 0.05 g/l ZnCl_2_, 0.03 g/l MnCl_2_·4H_2_O, 0.2 g/l CoCl_2_, 0.01 g/l CuCl_2_·2H_2_O, 0.02 g/l NiCl_2_·6H_2_O, 0.03 g/l (NH_4_)_2_MoO_4_·2H_2_O and 0.03 g/l FeSO_4_). For the experiment, M9 minimal salts medium with specified carbon sources and antibiotics was used. Antibiotics were used at the following concentrations: ampicillin (Amp), 150 μg/ml for *E. coli* and 500 μg/ml for *P. putida*; kanamycin (Km), 50 μg/ml; streptomycin (Sm), 50 μg/ml; gentamicin (Gm), 10 μg/ml; tetracycline (Tet), 50 μg/ml; and chloramphenicol (Chl), 34 μg/ml.

### Plate reader growth experiments

Strains were precultured overnight in LB with appropriate antibiotics. The next day, the cultures were centrifuged (1,500 g) and resuspended in M9 minimal medium. Growth assays were performed in Nunclon 96 Flat Bottom Transparent Polystyrene well plates (Thermo Fisher Scientific) containing 250 µl of M9 minimal medium supplemented with specified carbon sources (2 g/l unless stated otherwise) and antibiotics. Cultures were inoculated to an initial OD of 0,05 as measured in cuvettes with a standard 1 cm path length. The well-plate was covered with Breathe-Easy gas-permeable sealing membrane for microtiter plates (Diversified Biotech) and cultured in Infinite M Plex microplate reader (Tecan). Continuous shaking was set to orbital with an amplitude of 2,5 mm and each measurement was preceded by 10 s of linear shaking with an amplitude of 3 mm to avoid cell clumping. Fluorescence signal (bottom reading) and OD (OD_600nm_) were measured every half hour for 72 hours. Fluorescence gain was determined from undiluted positive controls and then set manually and maintained throughout the study (60 for GFP, 90 for mScarlet-I).

For recalculation of single strain fluorescence signal to single strain OD in cocultures, continuous calibration curves (fitted by combined linear or polynomial functions) of each strain grown on respective monosaccharides were prepared in each experiment. The wells containing calibration samples served simultaneously as a control dataset for the recalculation. Considering the delay of mScarlet maturation, the OD of CP-G was calculated as the OD of the consortium minus the OD of CP-X recounted from the reliable GFP signal.

### General cloning procedures

All plasmid constructs used and prepared in this study are listed in **Supplementary Table 2**. All plasmid constructs were built by standard restriction cloning or *in-vivo* cloning (Watson & Nafría) using chemocompetent *E. coli* CC118 or *E. coli* DH5α λπ (prepared in-lab with Mix & Go! *E. coli* Transformation Kit and Buffer Set from ZYMO Research). Plasmid DNA was isolated using E.Z.N.A. Plasmid DNA Mini Kit I (Omega Bio-Tek). DNA fragments were amplified by Q5 high-fidelity DNA polymerase according to the manufactureŕs instructions (New England BioLabs), purified using Monarch - PCR & DNA Cleanup Kit or DNA Gel Extraction Kit (New England BioLabs), and the purity and concentration were determined by NanoDrop 2000 (Thermo Fisher Scientific). Colony PCR for strain verification and plasmid build confirmation was performed in a 10 µl volume using 2x DreamTaq Green PCR Master Mix (Thermo Scientific) with oligonucleotide primers (0.5 µM each) and the addition of Q5 High GC Enhancer in case of *P. putida* templates. All primers used were purchased by Merck and are listed in the **Supplementary Table 3**. Their annealing temperatures were calculated using NEB Tm Calculator (New England BioLabs). Standard sequencing was performed by Eurofins Genomics or SEQme Czech Republic, whole plasmid sequencing by Plasmidsaurus (using Nanopore technology). Positive clones were stored in cryogenic stocks (20% v/v glycerol in LB) at −70°C.

### Strain engineering

Plasmids were inserted into *P. putida* by electroporation (2.5 kV, 25 µF, 200 Ω) in a 2 mm gap width cuvette (Thermo Fisher Scientific) using GenePulser XcellTM (Bio-Rad). The preparation of electrocompetent *P. putida* cells and the electroporation procedure were performed as described by Wirth et al., 2020. Cells were refreshed in LB medium after electroporation for 2 h.

For direct genome editing (deletions, insertions, site-specific mutations) we used a two-step method based on homologous recombination (HR) where integrative plasmids are built to redesign specific genomic loci (Martínez-García & De Lorenzo, 2012). The plasmids cointegrate into the genome by HR and a second recombination is induced by double-strand breaks upon *in vivo* cleavage by Sce-I homing endonuclease from *Saccharomyces cerevisiae*. The resolution of the HR event has a theoretical outcome of 50% in favour of the wild type and 50% in favour of the edited sequence. The protocol was optimised for *P. putida* by Martinéz-Garcia & Víctor de Lorenzo, 2012, where pEMG plasmid is used for cointegration and pSW-I for expression of Sce-I endonuclease. Upgraded versions of this protocol use pSNW plasmids for cointegration, and self-curing pQURE for Sce-I expression (Wirth et al., 2020; Volke et al., 2020). The glucokinase (PP_1011) deletion mutant Δ*glk* was built using pEMG. Insertion of degradation tags and deletion of N-terminal His tag was carried out according to the upgraded protocols. All genetic changes were checked by sequencing.

Tn7 insertions were carried out by mating. Four strains - *E. coli* CC118λπ pBG13/pBG13_mScarlet (bearing fluorescent protein genes as insertion cargo; donor strain), *E. coli* DH5α λπ pTnS-1 (strain leading transposase), *E. coli* pRK600 (helper strain) and a *P. putida* recipient strain - were mated by mixing equimolar amounts of cells in 10 mM MgSO_4_ and dropping the mix on an LB plate without antibiotics. The next day, a scoop of cells was resuspended in 10 mM MgSO_4_, diluted 100x, and 100 μl was spread on an M9 citrate (2 g/l) agar plate with 10 µg/ml gentamicin. The resulting *P. putida* transconjugants that exhibited fluorescence under blue light were selected for further work.

### Adaptive evolution of strain CP-X

The CP-X strain was grown in LB with kanamycin overnight. The next day, a 50 ml Erlenmeyer flask containing 6 ml of M9 minimal medium with kanamycin and 2 g/l of xylose was inoculated to a starting OD of 0,05. The culture was passaged after six days (OD reached 0,23), and a second time after 3 days (OD 1,5). The final culture was harvested after two more days of growth in fresh medium (OD > 1) and stored at −70°C in a cryogenic stock (20% v/v glycerol in LB). The parental strain was renamed CP-X(pre) and the evolved strain was named CP-X.

### Resting cell incubation for glycoside hydrolase activity testing

The conversion of disaccharide to monosaccharide by resting cells expressing β-glucosidase but unable to utilize glucose, and resting cells expressing β-xylosidase but unable to utilize xylose was monitored by HPLC. The test strains CP-X and CP-G and negative control strains without glycoside hydrolase enzymes (ID24 and ID18) were grown as night cultures and these were used to inoculate 10 ml of fresh LB medium with appropriate antibiotics and the strains were cultured for another 6 hours at 30 °C and shaking at 200 rpm in Shaking Incubator NB-205, N-BIOTEK. The culture sample (1 ml) of OD 1 was taken to prepare a cell sample and a supernatant sample by centrifugation. The cells were washed with pure M9 medium and resuspended in M9 medium with appropriate antibiotics. One more ml of the test strain culture (OD 1 or equivalent) was used to prepare a lysate sample by lysing the cells with B-PER (Thermo Scientific). A sample of the negative control strain cells prepared the same way as the cell sample of the test strain was used as the negative control without enzyme. All samples were diluted to an identical volume of 1,5 ml and supplemented with 2 g/l of disaccharide and appropriate antibiotics. Two antibiotics were used in supernatant samples to inhibit any additional cell growth (Km and Chl). The samples were incubated overnight at 30 °C with shaking at 200 rpm in Shaking Incubator NB-205 (N-BIOTEK). The next day, all samples were centrifuged and the resulting supernatants were prepared for HPLC analysis.

### HPLC analysis

The saccharide composition of growth experiment samples was analysed using the Agilent 1100 Series system (Agilent Technologies) equipped with a refractive index detector and Hi-Plex H, 7.7 x 300 mm, 8 µm HPLC column (Agilent Technologies). Collected bacterial cultures were centrifuged at 20,000 g for 10 min. The supernatant was filtered through 4 mm / 0.45 µm LUT Syringe Filters (Labstore), diluted 1:1 with 50 mM H_2_SO_4_ in degassed miliQ water to stop any hydrolytic activity, and stored at −20 °C in HPLC columns. Analysis was performed using the following conditions: mobile phase 5 mM H_2_SO_4_, mobile phase flow 0.5 ml/min, injection volume 20 µL, column temperature 65 °C, RI detector temperature 55 °C. Analyte concentrations were calculated from the area under the curve, related to respective analyte calibration curves prepared with pure chemicals purchased from Sigma-Aldrich (Merck).

### Whole genome sequencing

*P. putida* EM42-derived strains were cultured in LB medium at 30°C till mid-exponential phase. Cells were collected and enzymatically treated: 10 ml of bacterial culture in mid-exponential phase cultivated in LB at 30°C was centrifuged at 3,000 × g and 10°C for 10 min, washed with 5 ml of wash solution (10 mM Tris-HCl, 10 mM EDTA, 10 mM EGTA, 1 M NaCl of pH 7.5), and resuspended in Tris-EDTA (TE) buffer with achromopeptidase (1,000 U/ml; Sigma-Aldrich), lysozyme (5 mg/ml; Sigma-Aldrich), and RNase A (200 μg/ml; New England BioLabs) in a total volume of 500 μL, followed by incubation for 1 to 2 h at 37°C until lysis appeared. Then, 30 μL of 10% SDS and 5 μL of proteinase K (20 mg/ml; Sigma-Aldrich) were added, and the sample was incubated for 60 min at 50°C. Genomic DNA was extracted from cell lysate using the Genomic DNA Clean & Concentrator-25 kit (Zymo Research) according to the manufacturer’s instructions. For Oxford Nanopore sequencing, the library was prepared using the SQK-RBK004 Rapid Barcoding kit (Oxford Nanopore Technologies) according to the manufacturer’s instructions. The library was sequenced with a FLO-MIN106 flow cell (R9.4.1) in a MinION device controlled by MinKNOW software v.22.12.7, (Oxford Nanopore Technologies), which was also used for basecalling (super-accurate model with a minimum q-score threshold of 10), demultiplexing, and barcode trimming. Assembly of Nanopore reads was performed using Flye v.2.9.1 and Medaka consensus pipeline v1.7.2 (Oxford Nanopore). Genomic data was handled by Geneious Prime 2022.2.2 (Biomatters) and sequencing reads were mapped using the Geneious algorithm.

### SDS-PAGE

For each sample, 2 ml of cell culture of OD 1 was harvested by centrifugation at 4,000 g at 4°C for 10 minutes, and stored at −20 °C after decanting the supernatant. Cell-free extracts were prepared using B-PER reagent (Thermo Scientific). Centrifugation at 14,000 rpm (Hettich Universal 320R) and 4°C for 15 minutes separated the cell-free extracts from cell debris. The cell-free extracts were stored at −20 °C. Cell-free extract (4 μl) of each sample was combined with 2 μl of Sample loading buffer (5x; 1.2 ml Milli-Q H_2_O, 0.5 ml of 0,5M Tris-HCl pH 6.8, 0.8 ml glycerol, 0.8 ml 10% SDS, 0.2 ml β-mercaptoethanol, a pinch of Bromphenol blue) and boiled at 95°C for 5 minutes. The samples as well as 5 μl of Color Protein Marker II (NZYtech) were then loaded onto a 12% separating, 4% stacking SDS-PAGE gel and run for 35 minutes. The gel was stained by Quick Coomassie stain (Serva) for 1 h, destained in dH_2_O overnight, and scanned.

### Fluorescence microscopy

Cell samples were centrifuged (1,500g, 3 min), washed with M9 minimal medium, and fixed with 8% formaldehyde and 0.25% glutaraldehyde solution (1:1 v/v) for 30 minutes in Cellview cell culture slides (PS, 75/25 mm, glass bottom; Greiner). The fixed samples were washed with MilliQ water and covered with Vectashield Antifade Mounting Medium (Vector Laboratories). The cells were visualised using ZEISS Elyra 7 with lattice SIM mounted with the ZEISS objective Plan-Apochromat 63x/1,4 Oil DIC M27 and captured in pixel size 60 nm. The files were processed using SIM^2^ and images were prepared using orthogonal projection, all in ZEN Black software.

### Statistics

The number of repeated experiments or biological replicates is specified in figure and table legends. The mean values and corresponding standard deviations are presented.

## Acknowledgements

We are grateful to Dr. Tibor Botka from the Department of Experimental Biology, Faculty of Science, Masaryk University, for conducting the whole-genome sequencing of *P. putida* strains CP-Xpre and CP-X. We thank Alberto Sánchez-Pascuala for providing deletion plasmid pEMG _Δ*glk*HR. We also thank Victor de Lorenzo and Rodrigo Ledesma-Amaro for valuable discussions. This project was funded by the Czech Science Foundation (GAČR project registration number 22-12505S) and the Grant Agency of Masaryk University (GAMU MASH Junior MUNI/J/0003/2021) granted to P.D. and Brno Ph.D. Talent granted to B.B. This work was also supported by the ECCO (ERC-2021-COG-101044360) contract of the EU granted to A.G.M. and grants MULTI-SYSBIO (PID2020-117205GA-I00), BIOELECTRIC (CNS2022-135951), and MULTISYNBIO (PID2023-152470NB-I00) funded by MICIU/AEI/ 10.13039/501100011033.

## Contributions

**B.B.** conceptualized the work, performed experiments, cured data, and drafted the manuscript. **J.M.B.** developed the mathematical model, performed computational experiments, and cured data. **B.P.** and **B.G.** contributed by performing experiments and curing data. **A.G.M.** supervised **J.M.B. P.D.** conceptualized the work, secured resources, and supervised **B.B.**, **B.P.**, and **B.G.** All authors read, edited and approved the final manuscript.

## Data availability statement

The data that support the findings of this study are provided in the main text or available in the Supplementary Information of this article and from the corresponding author upon request. Raw data for all graphs displayed in the manuscript will be deposited in the Figshare repository. All sequencing data are stored under NCBI BioProject PRJNA1172556. Raw sequencing data have been deposited in the SRA database under accession numbers SRR31033117 (CP-Xpre) and SRR31033118 (CP-X). The whole-genome sequence of strain CP-X(pre) has been deposited in the GenBank database under accession number CP172071.

## Supplementary Information

### Supplementary tables

**Supplementary Table 1.** List of strains used and prepared in this study.

| Strain | Characteristic | Reference |
| --- | --- | --- |
| *E. coli* |  |  |
| CC118 | *araD139 Δ(ara-leu)7697 ΔlacX74 phoAΔ20 galE galK thi rpsE rpoB argEam recA* | Manoil & Beckwith, (1985) |
| DH5α λπ | DH5α lysogenized with λpir phage | Pablo Nikel Lab collection |
| DH5α λπ_pTNS-1 | Amp<sup>R</sup>, ori R6K, *tnsABCD*, strain with mobilisable helper plasmid | Choi et al. (2005) |
| HB101_pRK600 | Chl<sup>R</sup>, ori ColE1, tra+ mob+ of RK2, helper strain | Boyer & Dussoix, (1969) |
| *P. putida* EM42 |  |  |
| Δgcd | Chassis for the hexose eating strain | Dvořák & De Lorenzo (2018) |
| ID18 | Δgcd::mScarlet; Glycoside hydrolase<sup>-</sup> control for CP-G | This work |
| CP-G | Δgcd::mScarlet pSEVA2213_xyl43A; Hexose eater | This work |
| Δgcd::bglC | Tn5 inserted *bglC* gene from *T. fusca* (P<sub>em7</sub>, N-His *bglC*, Sm<sup>R</sup>); reverse strand in *xerD* (PP_1468) | Dvořák & De Lorenzo (2018) |
| Δgcd/Δglk::bglC | Δglk (PP_1011); chassis for a pentose eating strain unable to consume glucose | This work |
| ID24 | Δgcd/Δglk pSW-I; Glycoside hydrolase<sup>-</sup> control for CP-X | This work |
| CP-X(pre) | Δgcd/Δglk::bglC/msfGFP pSEVA2213_xylABE | This work |
| CP-X | Δgcd/Δglk::bglC/msfGFP pSEVA2213_xylABE; Pentose eater; CP-X(pre) adapted on xylose; ~275 kbp multiplication | This work |
| CP-X_ssrA | CP-X with *ssrA* degradation tagged *bglC* | This work |
| CP-X_ssrActrl | CP-X with *ssrActrl* degradation tagged *bglC* | This work |
| CP-X_6A | CP-X with 6A degradation tagged *bglC* | This work |
| CP-X-NHis400 | CP-X, *bglC* without N-His tag + synthetic RBS (400 a.u.) | This work |
| CP-X-NHis2500 | CP-X, *bglC* without N-His tag (2500 a.u.) | This work |
| CP-X-NHis2500_6A | CP-X_6A, *bglC* without N-His tag (2500 a.u.) | This work |

**Supplementary Table 2.** List of plasmids used and prepared in this study.

| Plasmid | Characteristics | Reference |
| --- | --- | --- |
| pSEVA2213 | Km<sup>R</sup>, ori RK2, P<sub>em7</sub> | Silva-Rocha *et al.* (2013) |
| pSEVA238 | Km<sup>R</sup>, ori pBBR1, XylS/Pm | Silva-Rocha *et al.* (2013) |
| pSEVA2213_*xylABE* | Km<sup>R</sup>, *xylABE* genes from *E. coli* | Dvořák & De Lorenzo (2018) |
| pSEVA2213_*xyl43ANHis* | Km<sup>R</sup>, *xyl43A* gene from *T. fusca* | This work |
| pEMG_Δ*glkHR* | Km<sup>R</sup>, ori R6K, HR spanning (~500 bp) upstream and downstream of glucokinase (PP_1011) in *P. putida* KT2440 genome | Alberto Sánchez-Pascuala, CNB-CSIC |
| pSW-I | Amp<sup>R</sup>, ori RK2, XylS/Pm→*I-sceI*, Conditional Sce-I endonuclease expression | SEVA collection |
| pMRE-Tn7-145 | Amp<sup>R</sup> Chl<sup>R</sup> Gm<sup>R</sup>, mScarlet-I | Schlechter *et al.* (2018) |
| pBG13 | Km<sup>R</sup> Gm<sup>R</sup>, ori R6K, Tn7L and Tn7R extremes, P<sub>em7</sub>, BCD2-msfGFP fusion | Zobel & Benedetti (2015) |
| pBG13_mScarlet | Km<sup>R</sup> Gm<sup>R</sup>, ori R6K, Tn7L and Tn7R extremes, P<sub>em7</sub>, BCD2-mScarlet-I fusion, mScarlet-I N'(MVSK > MIMGILSK) | This work |
| pQURE1L | Amp<sup>R</sup>, oriRK2, XylS/Pm→*trfA*, XylS/Pm→*I-sceI*, P<sub>14g</sub>, mCherry | Volke *et al.* (2020) |
| pSNW5 | Tet<sup>R</sup>, *traJ*, oriT, ori R6K, P<sub>14g</sub>, BCD2-msfGFP | Volke *et al.* (2020) |
| pSNW5_bglC-HR | pSNW5 with HR spanning (~500 bp) upstream and downstream of *bglC* 3' - end in CP-X genome | This work |
| pSNW5_bglC-HR_ssrA | pSNW5_bglC-HR with AANDENYALAA(*)<br>Encoding the ssrA degradation tag | This work |
| pSNW5_bglC-HR_ssrActrl | pSNW5_bglC-HR with AANDENYALDD(*)<br>Encoding the ssrActrl degradation tag | This work |
| pSNW5_bglC-HR_6A | pSNW5_bglC-HR with AAAAAA(*)<br>Encoding the 6A degradation tag | This work |
| pSNW5_bglC-HRw | pSNW5 with HR spanning (~500 bp) upstream and downstream of *bglC*, including the whole gene in CP-X genome | This work |
| pSNW5_bglC-HRw-NHis2500 | pSNW5_bglC-HRw without N-His (2500 a.u.) | This work |
| pSNW5_bglC-HRw-NHis400 | pSNW5_bglC-HRw without N-His,<br>+ synthetic RBS (400 a.u.) | This work |

Abbreviations. Antibiotic markers: Amp - ampicillin; Gm, Gentamicin; Chl, Chloramphenicol; Km - Kanamycin; Tet, Tetracycline; HR, homology region.

**Supplementary Table 3.**
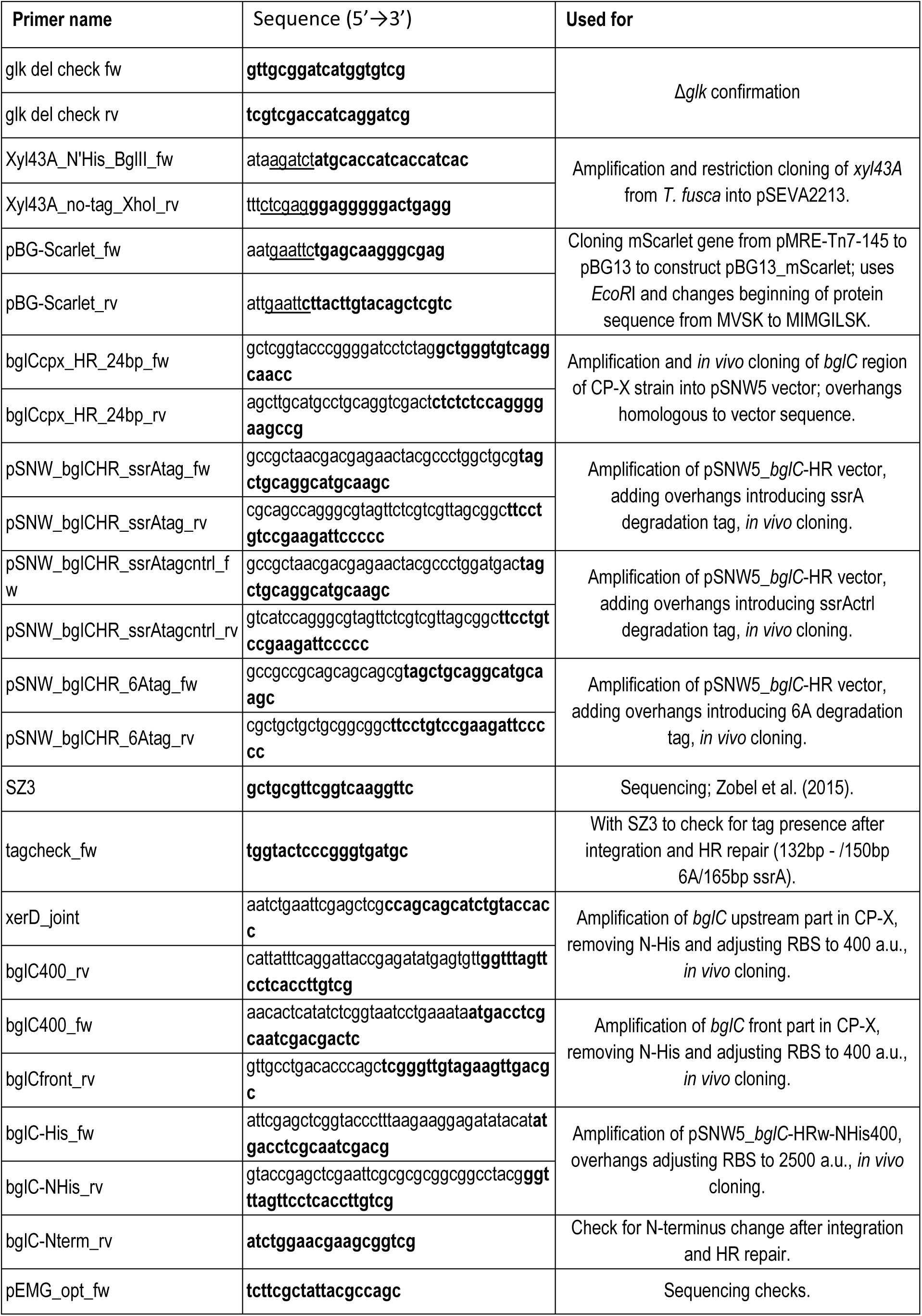
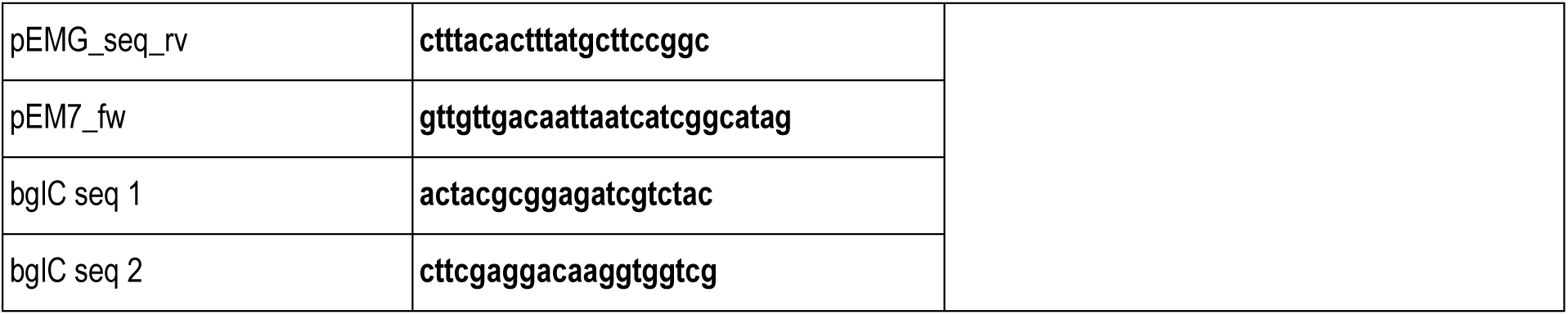
Primers used in this study. Annealing regions of the primers are highlighted in bold, and restriction sites are underlined.

**Supplementary Table 4.** Presence of a genomic multiplication in the chromosome of strain CP-X.

| Coverage |  |  |  |  |  |
| --- | --- | --- | --- | --- | --- |
|  | Chromosome without multiplied region (C) |  | Multiplied region (M) |  | M/C ratio |
| Strain | Nanopore reads | SD | Nanopore reads | SD |  |
| CP-X | 98.3 | 12.8 | 250.7 | 25.8 | 2.6 |
| CP-X(pre) | 77.4 | 11.5 | 77.2 | 6.9 | 1 |

Coverage was determined by the Geneious algorithm (Geneious Prime 2024.0.5). 275-kb region bordered by IS elements in pos. 2,220,489 - 2,495,683 bp in the reference genome of CP-X(pre).

### Supplementary figures

**Supplementary Figure 1.**
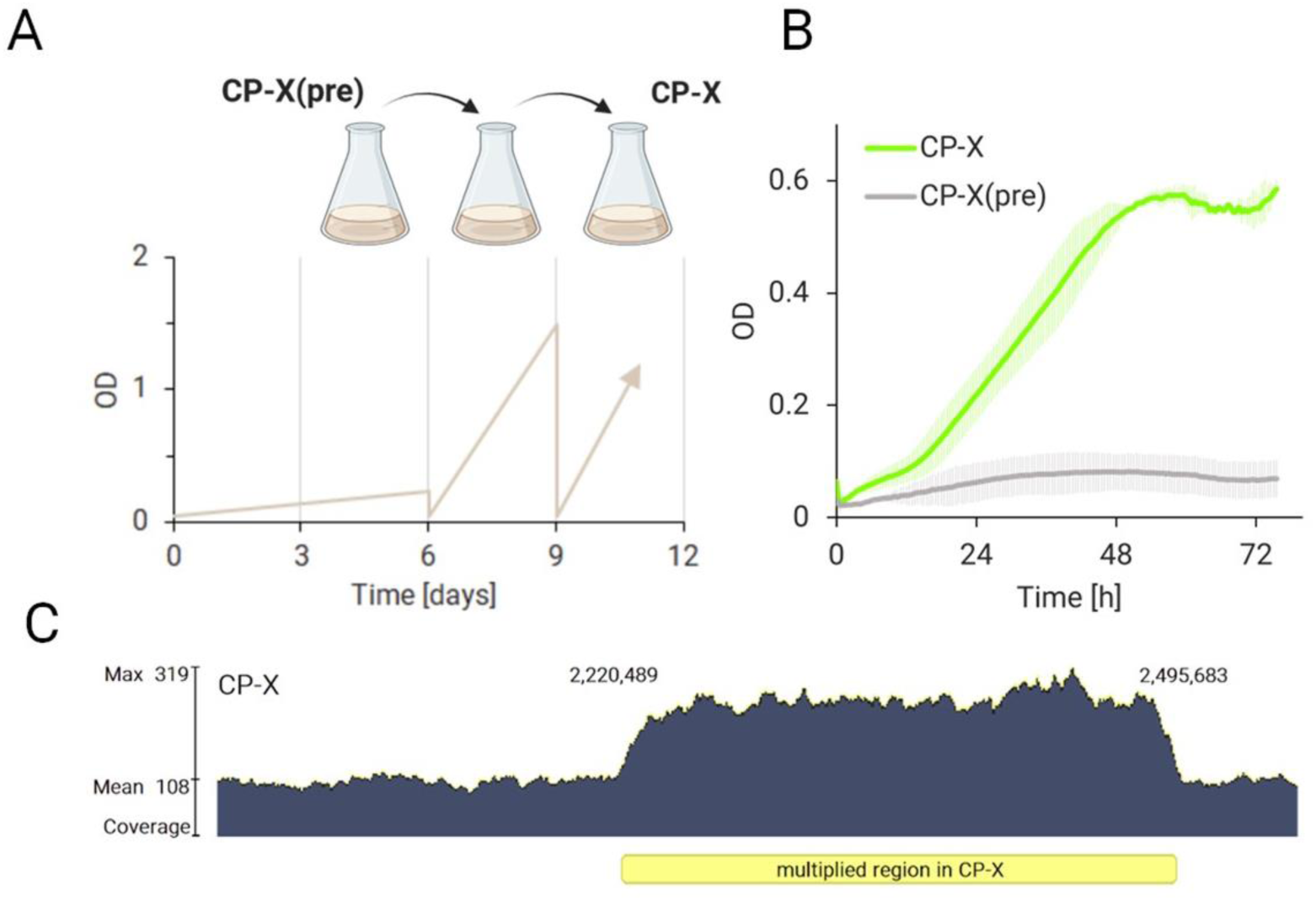
Xylose eater adaptation on xylose. (**A**) Reinoculation of the template strain CP-X(pre) in M9 minimal medium with 5 g/l of xylose as the sole carbon source led to an improved phenotype (CP-X). (**B**) Growth of both strains was compared in a 96-well plate cultivation on M9 medium with 2 g/l xylose. Data are shown as means ± SD from two (n = 2) biological replicates. (**C**) Sequencing revealed the emergence of a large genomic multiplication in the adapted strain CP-X.

**Supplementary Figure 2.**
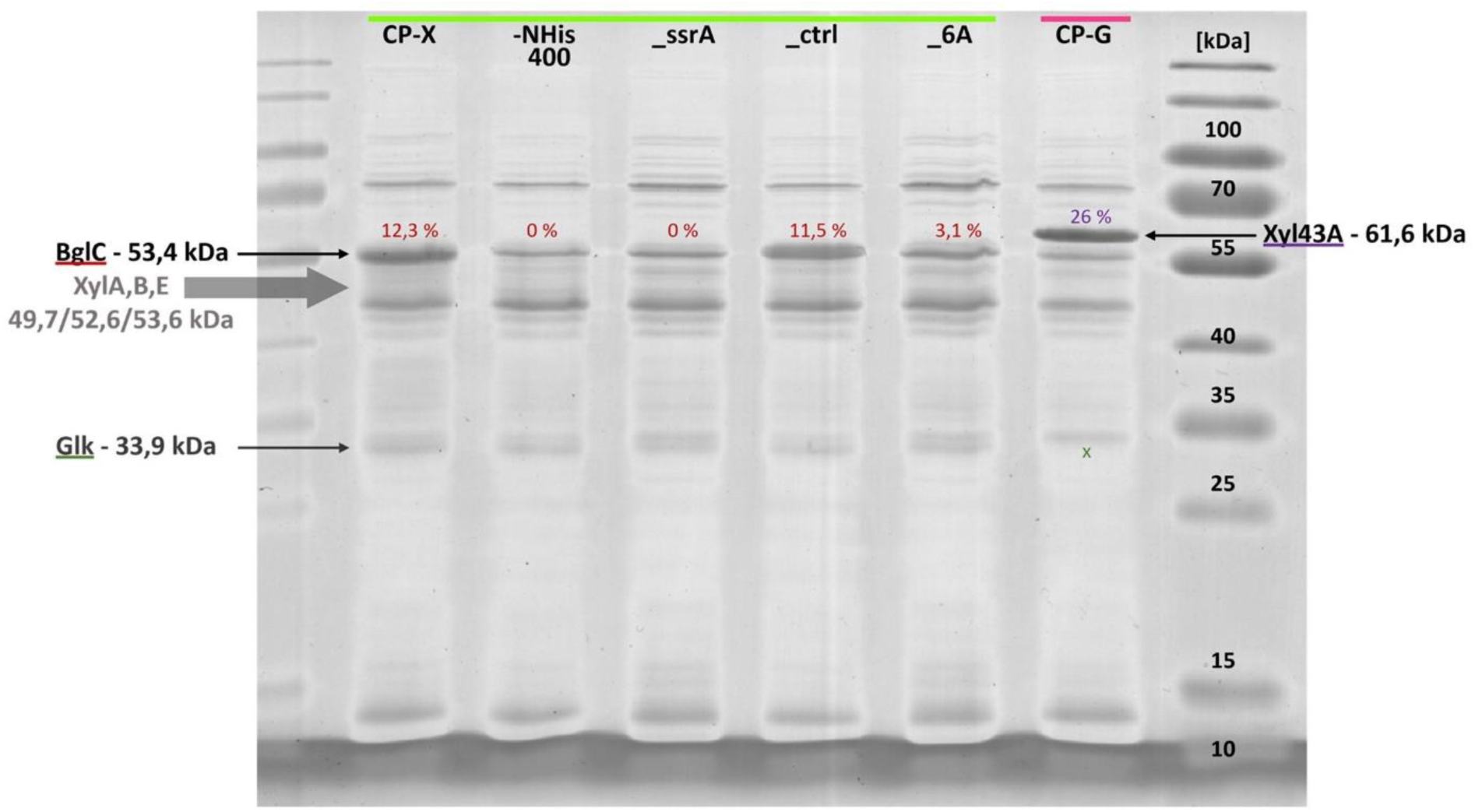
SDS-PAGE of consortium strain cell-free extracts. A 12% polyacrylamide (acrylamide and bis-acrylamide solution, 37.5:1) separating gel was used. Samples from the left: CP-X, CP-X-NHis400, CP-X_ssrA, CP-X_ssrActrl, CP-X_6A, CP-G. Protein abbreviations: BglC β-glucosidase, Glk glucokinase, XylA xylose isomerase, XylB xylulokinase, XylE xylose symporter, Xyl43A β-xylosidase.

**Supplementary Figure 3.**
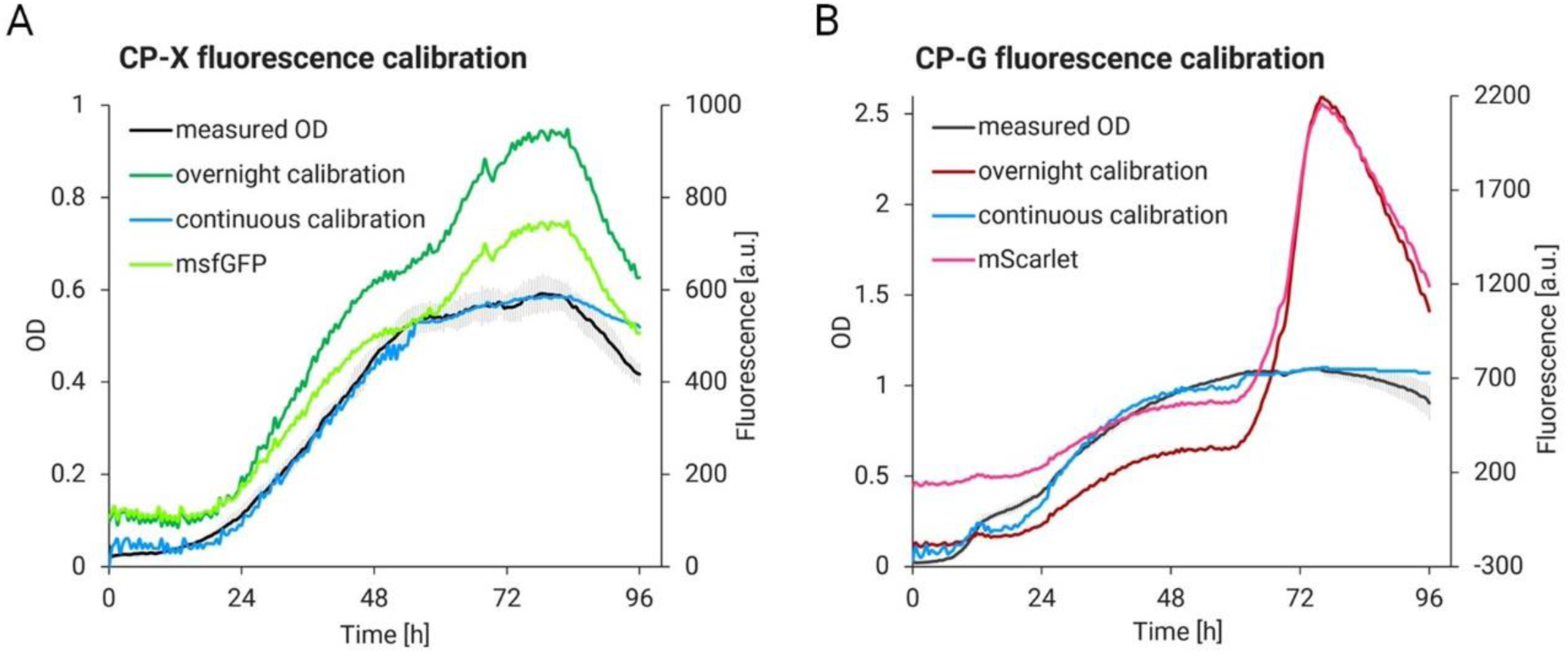
Comparison of fluorescence calibration options for the 96 well-plate cocultivation format. Overnight calibration was done by serial dilution of cells grown on rich LB medium to stationary phase. Continuous calibration was done by measuring fluorescence and OD in real-time during growth in M9 minimal medium with monosaccharides (2 g/l). Data are shown as means from four (n = 4) biological replicates. Error bars are omitted for clarity.

**Supplementary Figure 4.**
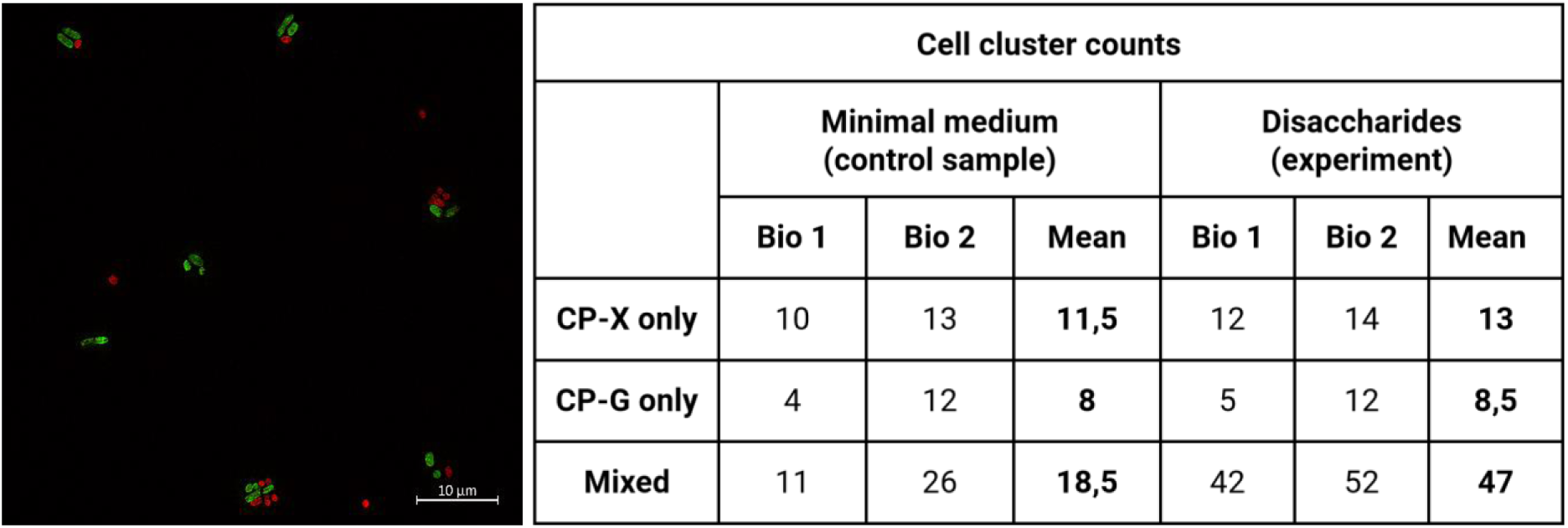
Cell cluster formation in the tightly linked consortium of CP-G and CP-X_ssrA. On the left, a micrograph of cell clusters recorded with ZEISS Elyra 7. Consortium of CP-G tagged by mScarlet and CP-X_ssrA tagged by msfGFP cooperating at growth on disaccharides. Photo taken at the end of cultivation (72 h). The table on the right reports the amounts of cell clusters formed during the growth in stress conditions (Minimal medium without carbon source) and cooperative growth on disaccharides. The experiment was conducted in two biological replicates (Bio, n = 2).

**Supplementary Figure 5.**
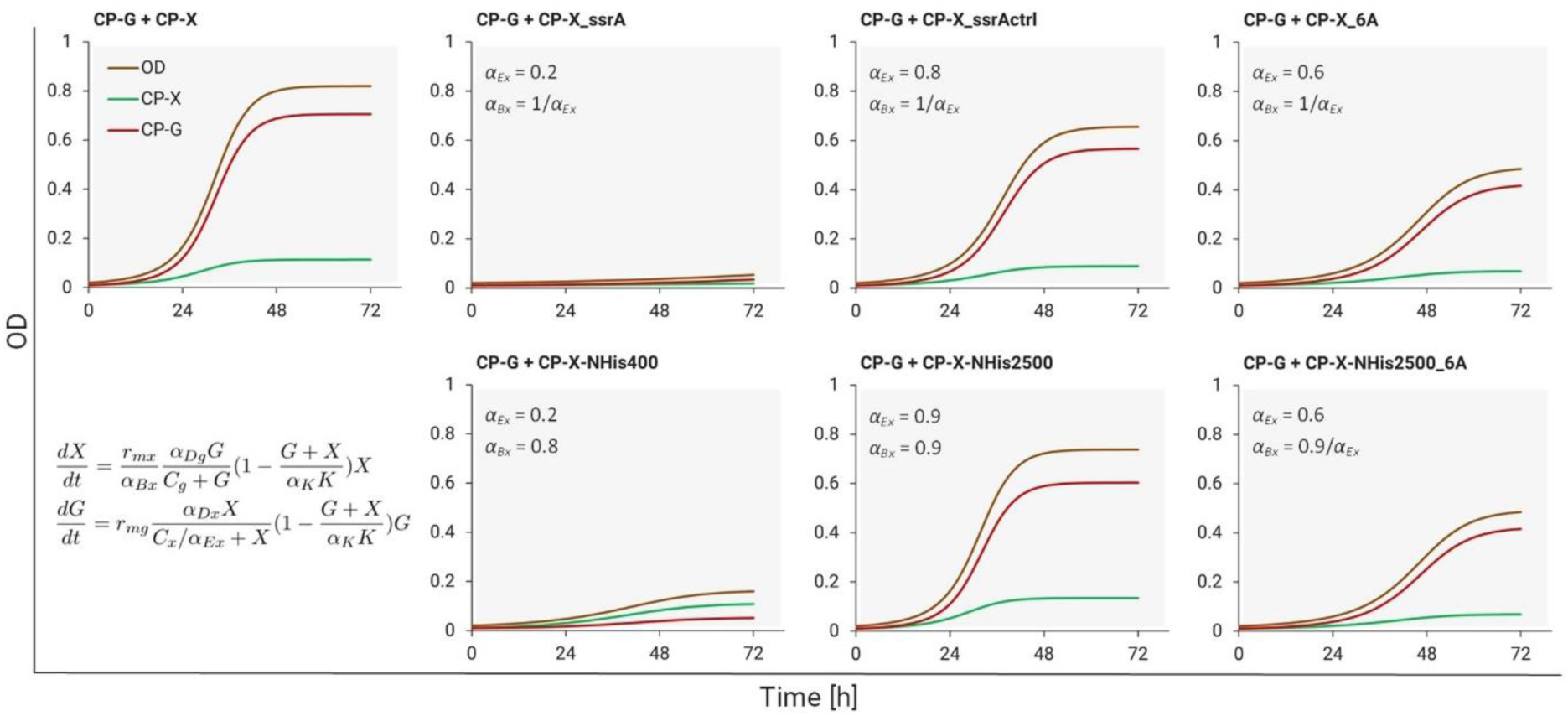
Mathematical model simulations of cooperating consortium strains with modulated levels of BglC expression and degradation. For the ctrl, 6A, ssrA, NHis400, NHis2500 and NHis2500_6A strains, the values of *α_Ex_* are 0.8, 0.6, 0.2, 0.2, 0.9 and 0.6 respectively. In ssrA, ctrl, and 6A strains, we assume the burden is inversely proportional to the amount of BglC, following the equation *α_Bx_* = 1/*α_Ex_*, to account for ATP-dependent BglC degradation. In NHis400 and NHis2500 strains, *α_Bx_* = 0.8 and 0.9, we assume *α_Bx_*<1 because the expression of BglC is lower, resulting in a decreased metabolic burden. For NHis2500_6A we use *α_Bx_* = 0.9/*α_Ex_* (*α_Ex_* = 0.6) to account for both the lower amount of BglC and the burden caused by BglC targeting to ATP-dependent proteasomes.

